# Mu-seq reveals previously undetected long-range contact patterns in the *Escherichia coli* nucleoid

**DOI:** 10.64898/2026.07.26.740832

**Authors:** Sonya Royzenblat, Khang Ho, Rucheng Diao, Rasika Harshey, Lydia Freddolino

## Abstract

The spatial organization of the *Escherichia coli* chromosome is a key determinant of its functional and regulatory landscape, but widely used chromatin conformation capture methods, such as Hi-C, provide an incomplete view of the underlying three-dimensional genome connectivity. Here, we introduce Mu-seq, a massively multiplexed Mu transposition-based approach that maps genome-wide DNA-DNA proximity in live cells without crosslinking. By distributing engineered, barcoded Mu prophages throughout the chromosome and tracking their replicative transposition events, Mu-seq generates dense, uniform sampling of physical contacts across the genome, enabling a direct, *in vivo* snapshot of chromosome architecture. Applying this method in *E. coli*, we constructed a contact dataset spanning over 200,000 unique loci, revealing interaction patterns not accessible to current crosslinking-based techniques.

Mu-seq findings can be broken down into three layers. First, it recapitulates polymer-like enrichment of short-range contacts. Second, contacts identified by Mu-seq are strongly shaped by chromatin state: transcriptionally active and accessible regions disproportionately contribute to long-range connectivity, whereas protein-occupied domains, including H-NS-enriched regions, are depleted of insertions and contacts. Third, beyond distance-dependent polymer behavior and chromatin accessibility, Mu-seq reveals specific long-range associations among functionally related loci, such as ribosomal operons. These findings suggest that bacterial chromosome organization can bring functionally related regions into shared three-dimensional neighborhoods even when they are widely separated along the linear genome.

Together, these findings support a layered model of nucleoid organization, in which transcriptionally active loops extend outward from a dense, protein-rich axial core. Mu-seq thus provides a powerful, complementary *in vivo* framework for probing bacterial chromosome architecture, uncovering regulatory and structural features of genome organization that are largely invisible to crosslinking-based approaches.

## Introduction

In *Escherichia coli*, and most prokaryotes, the genetic material resides within the cytoplasm alongside most cellular processes. DNA replication, transcription, and translation occur concurrently, within as little as 20 minutes in fast-growing cells (Royzenblat & Freddolino, 2024). While eukaryotic systems have long been known to rely on looping and long-range chromosomal contacts (e.g., enhancer-promoter interactions) to regulate gene expression over large genomic distances, the regulatory and structural roles of three-dimensional chromosome organization in bacteria have only recently come into focus.

Understanding chromosome conformation is critical for elucidating how regulatory proteins organize the genome and mediate interactions between linearly distant but spatially proximal loci. In bacteria, nucleoid-associated proteins (NAPs) can bridge and loop DNA to activate or repress specific genomic regions, contributing to diverse phenotypes including virulence-associated behaviors (Azam & Ishihama, 1999; Kotlajich et al., 2015),In addition, proteins can coat extended DNA regions, silencing transcription and, in some cases, forming phase-separated condensates (Amemiya et al., 2022b; Kahramanoglou et al., 2011; Lucchini et al., 2006; Rimsky et al., 2001). Together, these mechanisms establish chromosome architecture as a central regulatory layer in bacterial gene expression.

Yet, a key open question remains: to what extent do long-range chromosomal interactions in bacteria resemble the functional enhancer-like contacts seen in eukaryotes, and what regulatory roles do these interactions serve? The long-range interactions and spatial organization that comprise the 3D conformation of the chromosome are critical for efficient gene expression, replication, and repair, particularly in rapidly changing environments. While proteins such as sigma54 and NAPs can drive short- to intermediate-range looping (Dame, 2005; Ho et al., 2024; Nolivos et al., 2016; Reitzer & Magasanik, 1986; Su et al., 1990; Walker et al., 2020), it remains unclear whether truly long-range contacts - spanning hundreds of kilobases or more - play direct and specific roles in bacterial gene regulation. Resolving this question is essential for understanding complex phenotypes such as stress responses, persistence, and biofilm formation. A recent review has highlighted the importance of spatiotemporal chromosome organization, including NAPs, condensation, looping, and more (Royzenblat & Freddolino, 2024).

Microscopy has also revealed radial and subcellular organization within the bacterial nucleoid that may be difficult to capture using ensemble contact maps alone. H-NS, a major nucleoid-associated protein involved in transcriptional silencing and xenogeneic DNA repression, localizes to discrete clusters near the cell center and mediates long-range DNA-DNA interactions (Wang et al., 2011). In contrast, active transcriptional machinery and ribosomes show distinct spatial distributions. RNA polymerase can form organized bands and large foci during rapid growth, including clusters proposed to reflect intense ribosomal RNA transcription (Endesfelder et al., 2013). Ribosomes are strongly segregated from the nucleoid and enriched in surrounding cytoplasmic regions, consistent with models in which many transcripts diffuse outward from the nucleoid into ribosome-rich zones for translation (Bakshi et al., 2012, 2015). These observations support transertion-based and radial organization models in which transcriptionally active DNA may be preferentially exposed toward the nucleoid periphery, while protein-bound or transcriptionally silent DNA remains more centrally compacted (Bakshi et al., 2015). However, how these microscopy-defined spatial patterns correspond to genome-wide DNA-DNA contact architecture remains incompletely understood.

Over several decades, a wide range of approaches have been developed to investigate chromosome conformation. Early methods included fluorescence microscopy-based techniques and lambda recombination (Niki et al., 2000; Valens et al., 2004), followed by formaldehyde crosslinking-based sequencing methods such as 3C, Hi-C, and Micro-C (Gavrilov et al., 2025; Lioy et al., 2018). These approaches have dramatically improved our ability to resolve chromosome structure, but important discrepancies remain. For example, microscopy studies have consistently shown that six of the seven ribosomal RNA operons in *E. coli* cluster in three-dimensional space(Gaal et al., 2016), yet such clustering has not been detected in Hi-C or Micro-C datasets (Gavrilov et al., 2025; Lioy et al., 2018). These differences suggest that individual methods may preferentially capture distinct classes of interactions and highlight the need for complementary approaches.

To overcome limitations inherent to crosslinking and proximity ligation methods, we previously developed a Mu transposition–based strategy for detecting chromosome contacts (Walker et al., 2020). Mu is a bacteriophage that amplifies its genome through replicative transposition, inserting into distant chromosomal sites with little or no sequence specificity (Harshey, 2012, 2014; Mizuuchi, 1992). In the present application, barcoded Mu prophages are induced via a temperature-sensitive repressor, leading to expression of the transposition machinery. Importantly, transposition to a new site – detectable within 10 minutes post-induction – requires physical contact between donor and recipient DNA sites. Thus, mapping Mu start and insertion positions provides a readout of spatial proximity of DNA segments *in vivo*. In prior work, we demonstrated that this method recapitulates known contacts, including ribosomal operon clustering, and identified additional long-range interactions not detected by 3C-based approaches (which we hypothesized colocalize due to co-regulation), supporting a “small-world” model of chromosome organization (Lioy et al., 2018).

The original implementation of this method was relatively low throughput: Mu was inserted at 35 defined genomic locations, and each element was allowed to transpose once (Walker et al., 2020), equivalent to a series of 4C experiments. As a result, interactions were measured only from a limited set of starting points. To address this limitation, we developed Mu-seq, a massively multiplexed Mu-based approach that enables genome-wide sampling of transposition events. By distributing barcoded Mu elements broadly across the chromosome, Mu-seq achieves all-to-all contact mapping analogous in coverage to Hi-C but based on a fundamentally different physical principle: detecting DNA-DNA contacts *in vivo* without chemical crosslinking.

Using Mu-seq, we present a comprehensive analysis of *E. coli* chromosome conformation in live cells. Our results reveal that macrodomain boundaries are weak and permeable, consistent with a broadly interacting chromosomal polymer rather than rigidly partitioned spatial domains. We identify long-range interactions between functionally related loci that are not detected by crosslinking-based methods – ribosomal RNA operons preferentially contact one another, and additional clusters link distant operons involved in coordinated physiological systems, including stress response, envelope remodeling, and metabolic regulation. Finally, we demonstrate that chromatin accessibility, protein occupancy, and transcriptional activity collectively shape the interaction landscape. These findings support a layered model of nucleoid organization in which transcriptionally active loops extend outward from a dense, protein-occupied axial core, as initially proposed by Mäkelä and Sherratt (Mäkelä & Sherratt, 2020), and we further find evidence suggesting finer structure within the loops that distinguishes active from repressed chromatin regions. We also discuss possible future optimizations for better scaling of Mu-based contact mapping, particularly due to the limits imposed by barcode recovery at practical sequencing depths.

We begin by describing the construction and validation of the Mu-seq library, followed by analysis of global contact patterns, statistical modeling of long-range interactions, and integration with chromatin state, supercoiling, and transcriptional data. Together, our work establishes Mu-seq as a powerful and complementary approach for probing bacterial chromosome architecture, providing a direct *in vivo* perspective and revealing features of chromosomal connectivity that are not captured by existing crosslinking-based methods.

## Results

### Mu-Seq Library Construction and Validation

Mu-Seq (Ho et al., 2024) maps physical proximity across the *E. coli* chromosome by tracking replicative Mu transposition events in live cells. Based on our earlier work, we understood that Mu was able to transpose efficiently to sample the entire chromosome (Walker et al., 2020), but the throughput was limited. We thus designed a system in which a unique barcode on each of the prophages enabled us to sample and track each Mu prophage transposition (Figure 1A), providing a much more diverse starting and final library of Mu integrants than the initial 35 prophages described previously.

**Figure 1:**
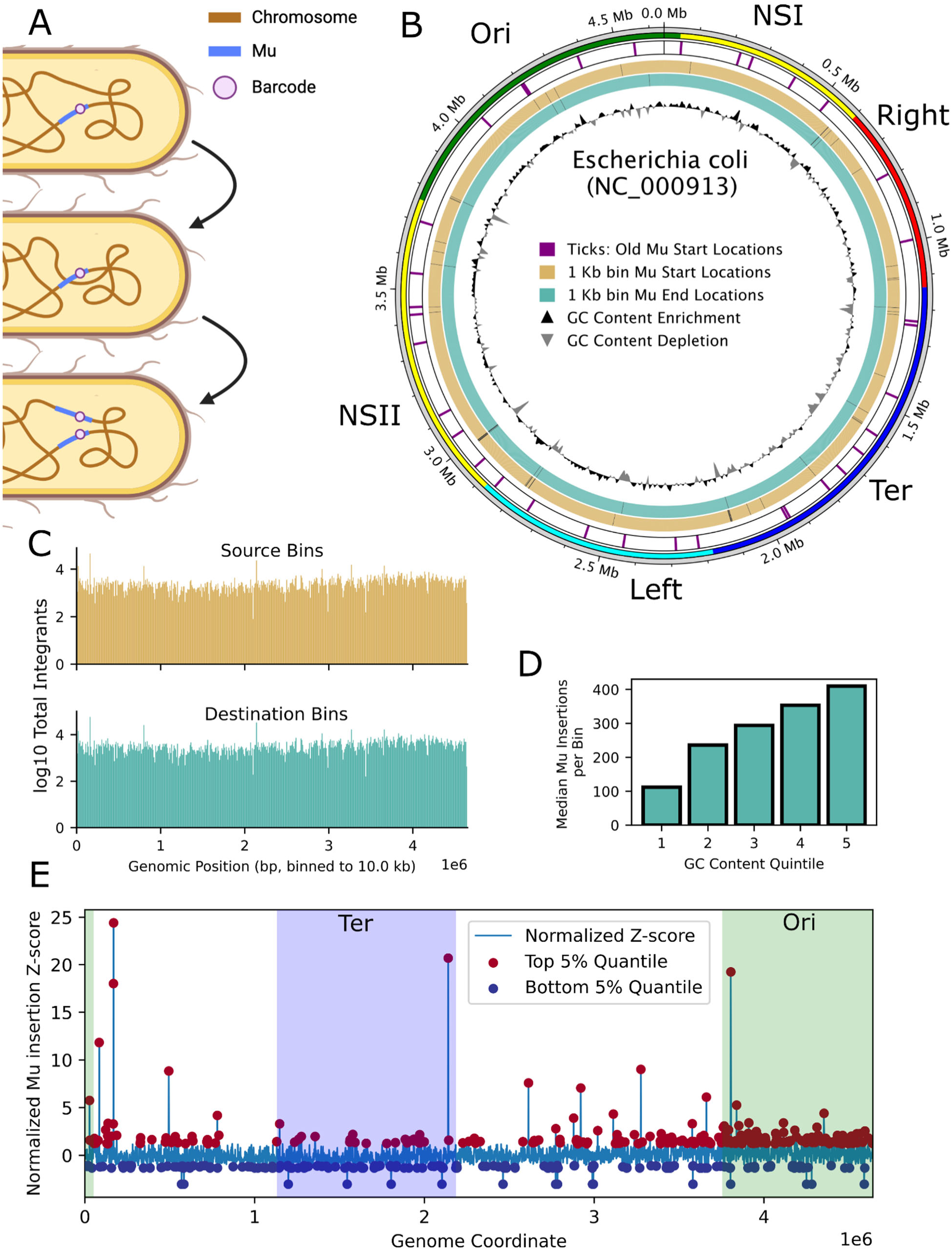
Construction and validation of high-throughput Mu transposition library. **A)** Schematic of Mu-based chromosome conformation capture method, illustrating barcode movement via Mu transposition to physically proximal regions. See Methods and (Ho et al., 2024) for additional details. **B)** Moving inward: macrodomains (Valens et al., 2004) are labeled at the periphery; purple ticks mark the 35 start sites from Walker et al., 2020 (Walker et al., 2020); orange and turquoise ticks mark source (uninduced) and destination (induced) library insertion locations, respectively, binned to 1 kb; black ticks indicate bins with no detected Mu insertion; the innermost ring shows GC content by region. Note, coverage is very dense and so individual ticks are not resolvable, except for the bins absent of Mu insertions. **C)** Distribution of Mu integrations densities from source (uninduced, orange) and destination (induced, turquoise) libraries, binned to 1 kb. **D)** Median Mu insertions per 1 kb bin across GC content quintiles, showing a positive relationship between GC content and insertion frequency. **E)** Normalized Mu insertion enrichment and depletion (Z-score) across the circular *E. coli* genome in 1 kb bins. Top and bottom 5% outliers are highlighted; shaded regions indicate Ori (green) and Ter (blue) macrodomains (Valens et al., 2004).

To construct barcoded donor libraries, we integrated the unique barcode into each Mu prophage via the lambda integrase system near the right end of Mu (MuR) (Ho et al., 2024). This barcoded prophage library was induced to generate particles carrying each barcode, which were then used to infect fresh cells – creating the initial donor library. Transposition was triggered using a temperature-sensitive repressor to express Mu transposition machinery, permitting a single round of replicative transposition without activating the lytic cycle or causing cell lysis. This controlled approach enables precise labeling of new insertion sites across the chromosome, capturing a high-resolution snapshot of chromosome conformation in intact cells.

Each prophage in the initial lysogen library defines a “source” site; each new integration event defines a “destination” site physically proximal to the source at the moment of transposition. Each Mu element inserts into genomic loci that are spatially proximal to its source site in three-dimensional space, thereby recording local DNA–DNA contacts as transposition events. Libraries were processed as described in Ho et al., 2024 (see Methods for additional details). We detected nearly 200,000 locations in our source library and approximately 230,000 in our destination library, tracking 119,106 unique barcodes in total. The library exhibited substantial diversity and broad genomic coverage (Figure 1B–C, Supplementary Figure 1), with most barcodes occurring near the mean frequency. A minority of barcodes occurred at substantially higher frequencies, but rank-abundance plots and barcode-by-location scatterplots show no evidence of unintended bottlenecks or excessive redundancy of barcodes (Supplementary Figure 1A–D).

Plotting source and destination locations binned at 1 kb (Figure 1B) illustrates the dramatic increase in coverage relative to our prior dataset (Walker et al., 2020): from 35 unique source sites to 198,294 source locations (purple) and 234,059 destination locations (turquoise).

The distribution of barcode integration densities was right-skewed, with most barcodes detected at modest frequencies and a minority occupying “hyperactive” sites with many unique barcodes (Figure 1C, Supplementary 1A-B). These high-abundance regions qualitatively align with the boundaries of the Ori/Ter organizational pattern (Figure 1E), defining their borders and likely correspond to regions of elevated chromatin accessibility.

To test whether there was a macrodomain preference for insertions, we normalized Mu insertion rates genome-wide and applied Mann–Whitney tests to the Ori and Ter macrodomains. After normalizing for read depth, the median coverage-normalized insertion rate in the Ori region (1.92 insertions per barcode per 1 kb bin) was modestly but significantly higher than the genome-wide median (1.69; U = 2,238,789.50, p = p=1.04e-44) while the Ter median was slightly lower (1.57; U = 1,061,593.00, p=2.65e-35). This suggests a genuine, if modest, preference for insertion near the origin beyond simple copy-number effects.

Mu exhibits a slight sequence preference for regions containing the CGG motif (Manna et al., 2005). Consistent with this known selectivity, binning the *E. coli* genome into 1 kb intervals grouped by GC content quintile reveals that the median number of Mu insertions per bin increases with GC content (Figure 1D). Despite this preference, we highlight in Figure 1C that our library captures transposition events with high density and genome-wide uniformity.

To assign transposition events, we cross-referenced the uninduced and induced libraries, identifying barcodes whose genomic location shifted between conditions. After stringent filtering to exclude ambiguous barcodes (those found at multiple locations in the uninduced library), we detected 2,589 unique barcodes and 4,476 barcode-location pairs that could be confidently linked across uninduced and induced samples across all replicates. This number was surprisingly limited given our ∼100,000-barcode input and the extensive sequencing depth employed (1.2 billion reads in a first run and 150 million in a second); we address this challenge further in the Discussion.

To maximize the number of usable contacts, we developed a complementary “one-sample” analysis. Because Mu undergoes replicative transposition, any barcode detected at two locations in the induced library must reflect a transposition event between those sites - one position being the source, one the destination - with barcode collision being exceedingly unlikely given library diversity. We therefore treated such pairs as non-directional contacts (with 119,106 unique barcodes distributed across a 4.6 Mb genome, the probability that two independent Mu elements share the same barcode and happen to insert near the same destination site is vanishingly small (∼10⁻⁶ per event), making coincidental barcode collision an implausible explanation for any observed two-location barcode). This is distinct from the “two-sample” analysis, which required barcodes to be assignable in both uninduced and induced libraries. The one-sample approach added 36,900 unique barcode-location pairs to our analysis.

Replicate-level quality control supported use of the consolidated dataset: binned insertion profiles were significantly correlated across replicates, with Spearman correlations of ρ = 0.41–0.65 for source-site abundances and ρ = 0.80–0.94 for destination sites across all pairwise comparisons (Supplementary Figure 1C–D). In total, our consolidated contact library contains 18,573 unique locations, 38,551 transposition hops, and 34,565 self-transitions (hops within 200 bp of the source site), with a mean of approximately 2 transitions per barcode (Supplementary Table 1).

Together, these results establish Mu-seq as a high-complexity, genome-wide contact mapping resource. We next turned to the broader organizational features of the *E. coli* chromosome revealed by this dataset.

### Global Patterns in Mu-Detected Chromosome Contacts

To explore global chromosome conformation patterns and assess higher-order structures detectable by Mu, we aggregated Mu insertion pairs into a 46 kb-binned sequential component normalized (SCN) contact matrix (Figure 2A) – that is, the frequency of contacts measured between bins, controlling for the abundances of DNA from each bin (see Methods for details). The heatmap reveals modest structured patterns along the diagonal reminiscent of chromosome interaction domain (CID)-like regions, though without the distinct macrodomain structures seen in Hi-C maps. We do, however, detect elevated off-diagonal contacts indicative of specific long-range interactions (discussed below). Ori- and Ter-proximal regions behave distinctly: Ori displays elevated interaction propensity while Ter shows reduced contact signal consistent with relative isolation. Broader contact patterns are therefore most apparent in non-Ori/Ter regions (Figure 2A-B, Supplementary Figure 2A, Supplementary Table 2). Notably, certain bins acting as Mu donors show significantly more contacts genome-wide than others, suggesting that some loci are less locally constrained and more permissive to distal Mu-mediated contacts; this is quantified further below.

**Figure 2.**
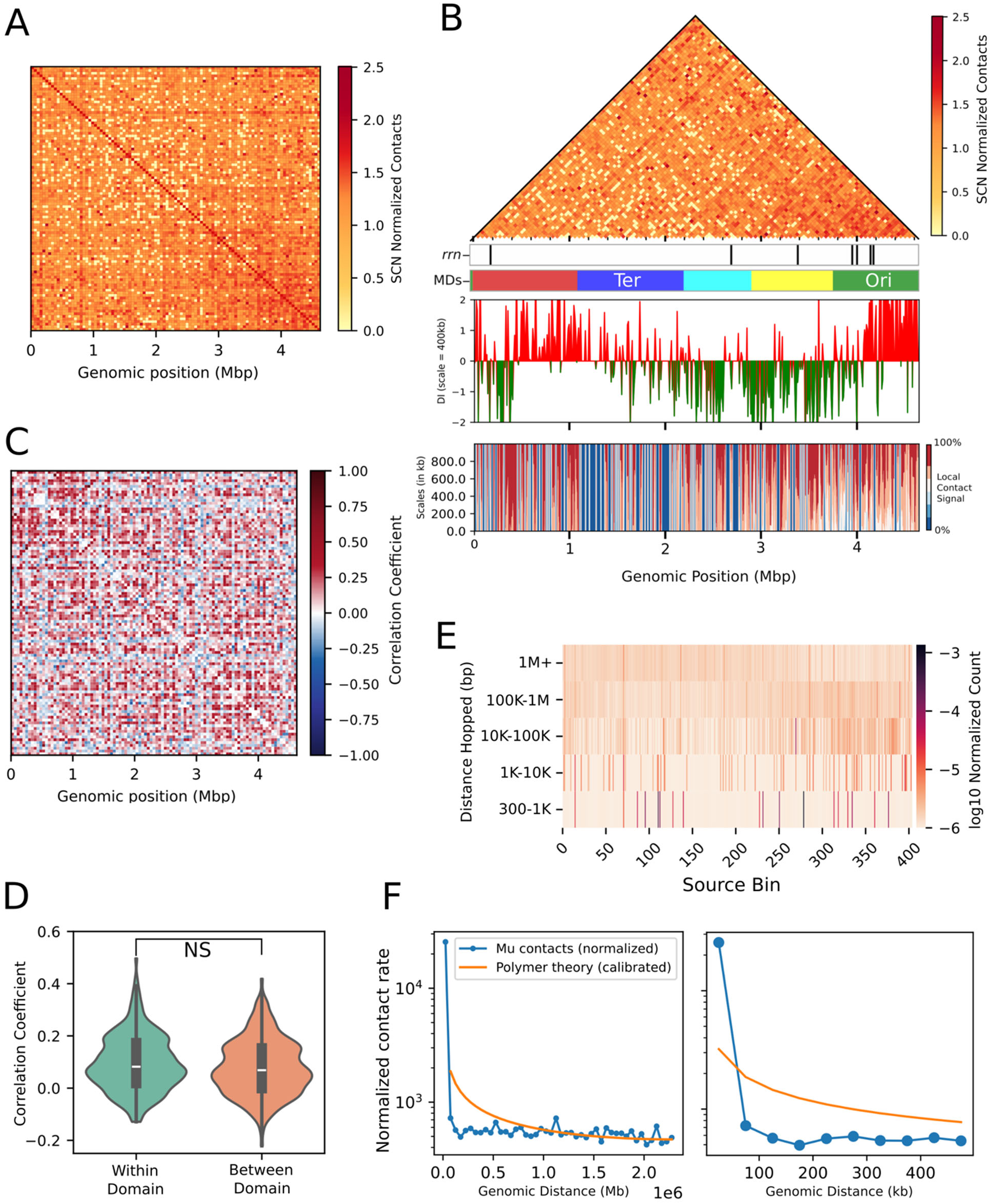
Genome-wide Mu contact patterns and chromosome organization. **A - B)** SCN contact maps (46 kb bins) reveal broad spatial interaction patterns with only modest domain-like insulation. Panel B additionally annotates ribosomal operon locations (rrn) and macrodomain (MD) boundaries for reference. Middle: directionality index (400 kb window) depicting upstream/downstream contact bias; red and green indicate opposite directional biases. Bottom: scale-dependent local contact signal, computed as summed SCN-normalized contacts within increasing windows around each genomic bin; warmer colors indicate higher local contact enrichment at that scale. **C)** Pearson correlation matrix of distance-normalized Mu contact frequencies across 46 kb bins. Off-diagonal clusters identify regions with similar contact profiles beyond linear proximity; overall compartmentalization is weak. **D)** Violin plot comparing within-macrodomain versus between-macrodomain correlation coefficients from panel C. Macrodomain insulation is detectable only for proximal bin pairs and was not found statistically significant by a two-sided permutation test comparing the difference in mean correlations. **E)** Heatmap of Mu transposition hop frequencies across binned genomic distances, normalized by opportunity and shown on a log₁₀ scale. The y-axis represents distance hopped; the x-axis denotes source location (binned to 11.5 kb). Opportunity is higher for longer-distance coordinate pairs. Sampling is uniform across all genomic bins, with no regional bias for short- or long-range contacts. **F)** Distance decay of Mu hops (10 kb bins) as a function of genomic distance, normalized per opportunity (observed insertions divided by possible locus pairs at each distance). A calibrated polymer scaling law (1/√distance) is overlaid. The lower panel zooms to the first 500 kb. The observed decay is consistent with polymer model predictions.

The directionality index (DI) (Lioy et al., 2018) revealed sharp transitions in upstream versus downstream contact bias throughout the chromosome (Figure 2B, middle), rather than the broad macrodomain-scale plateaus one might expect. These transitions were distributed across much of the chromosome, with Ori and Ter again behaving distinctly. Importantly, these sharp DI peaks are not driven by local transcriptional propensity (Scholz et al., 2019, 2022) nor correlated with transcript levels (Supplemental Figure 2B-C). The scalogram (Figure 2B, bottom) similarly revealed heterogeneous, position-specific local contact enrichment across scales, rather than broad continuous macrodomain-like blocks, consistent with an organizational landscape dominated by narrow frontiers and heterogeneous local compaction. A frontier signal map (Supplemental Figure 2C) – which highlights genomic positions where contact patterns change abruptly between adjacent bins– highlights these transitions genome-wide: their borders align closely with known CID boundaries, yet we do not observe the strong within-domain contact enrichment and between-domain depletion expected under a classical macrodomain model. The prominent boundary signals in our frontier analysis therefore suggest that whatever factors define macrodomain boundaries in Hi-C do not prevent Mu transposition across them.

### Correlation analysis detects weak insulation

We were able to identify an additional layer of structural organization by computing pairwise Pearson correlations of interaction profiles across 46 kb bins, thus providing a ‘guilt-by-association’ measure of which regions showed similar overall contact patterns (Figure 2C). Neighboring bins show broadly similar interaction profiles, as reflected by high correlations along the diagonal, particularly near Ori. Beyond the diagonal, discrete clusters of long-range interactions are visible as red blocks surrounded by blue perimeters, suggesting preferred inter-domain contacts or higher-order folding.

Following macrodomain definitions from Valens et. al. (Valens et al., 2004), we quantified within-versus between-domain correlation coefficients. Proximal within-domain bin pairs (≤10 bins apart) had a modestly higher mean correlation (0.101) than between-domain pairs (0.093), but this difference was not found statistically significant by a two-sided permutation test comparing the difference in mean correlations (p = 0.0831; Figure 2D). Distal within-domain pairs (>10 bins apart) showed a mean correlation of 0.086, also not significantly different from between-domain pairs (p = 0.3777, permutation test). Isolated blocks of self-correlation near Ori and Ter, alongside broadly distributed positive and negative correlations across domain boundaries, indicate a complex and weakly insulated chromosomal landscape. Taken together, any macrodomain insulation detectable by Mu-seq is modest and confined to near-diagonal, local interactions.

### Quantitative Analysis of Distance Decay and Spatial Organization

To assess the range and genomic distribution of Mu transposition contacts, we constructed a heatmap of opportunity-normalized, log-transformed contact frequencies binned by both genomic position and hop distance (Figure 2E). Higher-intensity signals reflect bins with increased Mu insertion rates after controlling for the number of possible source-destination pairs at each distance.

Short-range hops were strongly enriched relative to longer-range hops: those spanning <100 kb had a mean observed/expected enrichment of 12.23, whereas hops ≥100 kb had a mean enrichment of 0.50 (difference = −11.73, p = 0.0016, permutation test; Supplementary Tables 3-4). This preference for nearby loci is consistent with polymer theory predictions, in which contact frequency scales with the inverse square root of linear genomic distance (Figure 2F). Importantly, however, when normalized for sampling opportunity, Mu transposition can access the full range of genomic distances - in contrast to Hi-C, where distance-decay normalization still reflects a strong bias toward short-range contacts, likely introduced by crosslinking during sample preparation. This ability to sample contacts at any linear distance is a key advantage of the Mu-seq approach. We also note that a strong local-hop bias is similarly observed at lower induction levels in the lambda recombination experiments that originally established macrodomain definitions (Figure 2C of Valens et al., 2004).

The weak macrodomain insulation seen in our correlation matrices (Figure 2C-D) is consistent with a broadly distributed contact architecture, in which closely spaced bins show only marginally increased cohesion within domains and strong compartmentalization is absent (Walker et al., 2020). Taken together, these findings recapitulate the large-scale observations from our earlier low-throughput study while extending them to longer-range contacts not accessible to 3C-based methods.

### Statistical Modeling of Chromosomal Interactions

While these global patterns establish the broad organizational logic of the chromosome, they do not resolve whether specific distant loci interact preferentially beyond what polymer behavior and chromatin state would predict. To identify such interactions systematically, we applied a Poisson regression model with LASSO regularization, modeling observed contact counts between genomic bins as a function of inter-bin distance, DNA abundance, and replicate (see Methods). The fitted pairwise interaction strengths between bins - after controlling for these factors - are referred to as interaction coefficients; elevated coefficients between specific bin pairs indicate enrichment for long-range chromosomal contacts. We refer to the fitted interaction strengths between bins (after controlling for the other factors enumerated here) as the interaction coefficients; specific pairwise interactions between bins (with high interaction coefficients) may indicate enrichment for long-range chromosomal contacts.

The interaction coefficients recapitulate the key conclusions from our distance-decay analysis and from our prior low-throughput Mu experiments (Walker et al., 2020): contacts remain detectable across the full range of linear distances, with no sharp discontinuities or strongly isolated interaction blocks (Figure 3A). While contact frequency decays weakly with distance, consistent with polymer behavior, chromosomal regions remain broadly interactive. These data are consistent with a “small-world” model of *E. coli* chromosome organization, in which virtually any region can interact with any other without confinement to local domains. Nevertheless, we do observe distinct clusters of distal genomic bins that interact preferentially with one another, pointing to specific long-range associations among functionally related loci.

**Figure 3.**
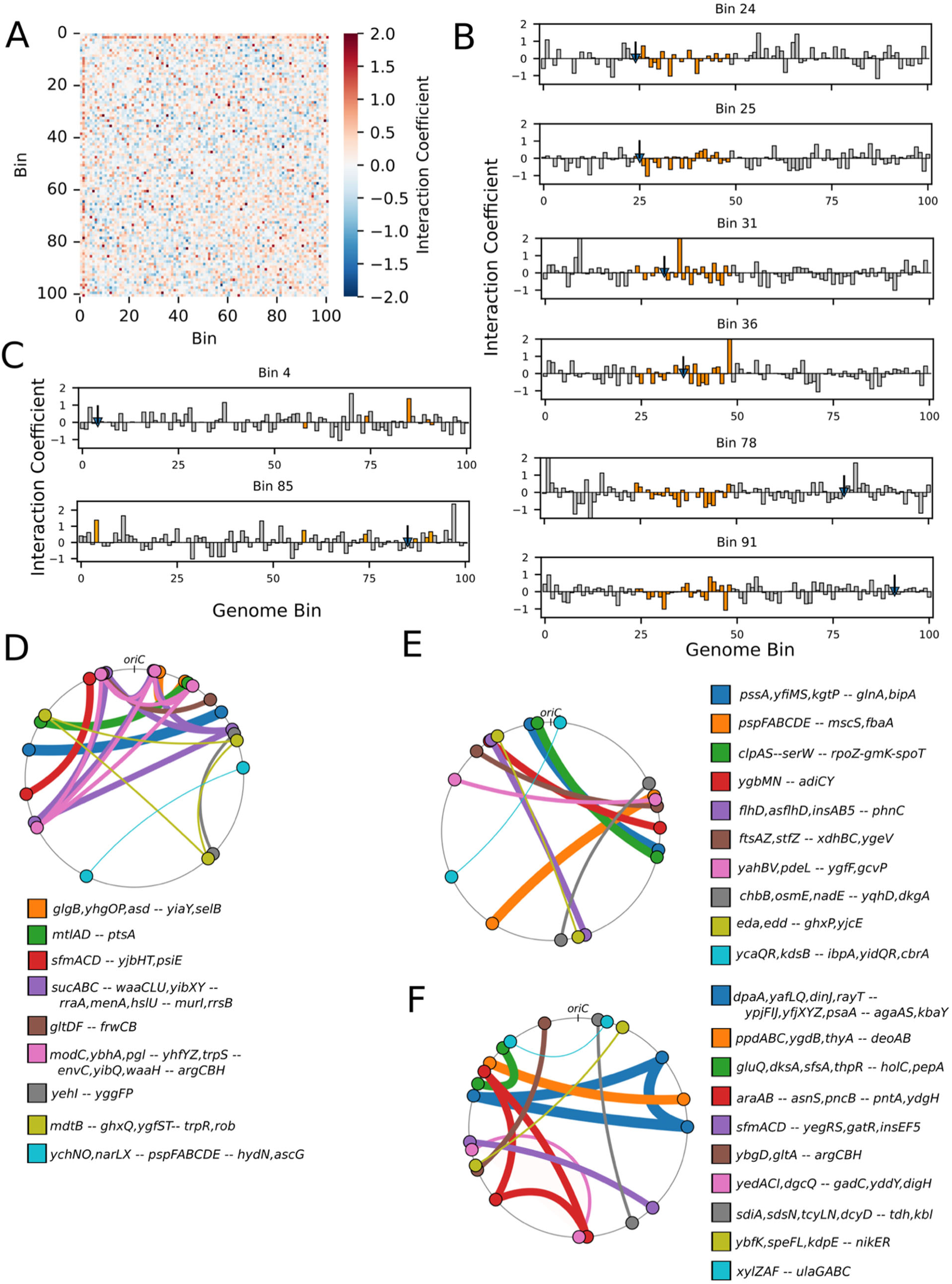
Modeling Mu-derived chromosomal contacts and long-range clustering. **A)** Poisson regression heatmaps of interaction coefficients between genomic bins, capturing both local and distant chromosomal contacts. **B)** One-dimensional donor/recipient contact profiles for example bins (24, 25, 31, 26, 78, and 91), enlarged from Panel A. Source bin marked with a blue arrow. Bins within the Ter macrodomain are highlighted in orange. **C)** One-dimensional donor/recipient contact profiles for bins 4 (*rrnG*) and 85 (*rrnC*), enlarged from Panel A. The source bin is marked with a blue triangle. Bins containing ribosomal RNA operons are highlighted in orange. **D-E)** Circos cluster plots visualizing significant long-range contacts between operons identified by the bin-shifting pipeline, with arcs connecting bins that show robust interaction coefficients across coordinate shifts, and arc weights corresponding to relative Spearman correlation values, with the strongest correlated clusters containing the thickest arc width.

Examining bin-specific interaction coefficients confirms established trends for the Ter macrodomain (bins 24–48), which consistently display lower interaction coefficients, indicative of reduced colocalization and relative chromosomal insulation (Figure 3B). Bins at the Ter boundary (e.g., bin 24) show somewhat elevated interaction coefficients, whereas bins just inside the boundary (e.g., bin 25) exhibit notably diffused interaction profiles, and bins far outside the Ter (e.g., bins 78 and 91) interact broadly but show decreased correlation specifically with Ter bins. This pattern supports the notion that the Ter region presents a relative barrier to Mu penetration, potentially due to protein-mediated insulation (e.g., MatP binding (Mercier et al., 2008; Walker & Harshey, 2020)) or higher local compaction of chromatin structure – limiting Mu accessibility both within and across the Ter boundary. Interestingly, when Mu is observed starting in a bin well in the middle of the Ter (e.g., bins 31 and 36) (Figure 3B), we see not only relatively decreased interactions with the majority of bins outside the Ter, but also negative interaction coefficients with bins even within the Ter. One interpretation is that Ter-proximal chromatin is comparatively compact or constrained, reducing Mu accessibility to both within Ter and between Ter and adjacent regions. The boundary is nonetheless weak: contacts between Ter and non-Ter bins persist, and the depleted contact pattern extends beyond the canonical Ter boundary definition.

Interestingly, bins containing ribosomal operons exhibit preferentially elevated interaction coefficients with other *rrn*-containing bins (Figure 3C). For example, Mu transposition initiated from bin 85 (*rrn*C) or bin 4 (*rrn*G) yields interaction profiles with consistently positive coefficients at other *rrn* bins (marked “o”). This finding echoes results from both previous Mu-based studies (Walker et al., 2020) and microscopy data reporting colocalization of 6/7 ribosomal operons (Gaal et al., 2016). One apparent discrepancy persists: transposition from bin 4 (*rrn*G) yields a negative interaction coefficient for bin 85 (*rrn*C), whereas transposition initiated at bin 85 shows positive coefficients with all other *rrn* bins. A possible explanation is that the *rrn*C locus resides in a generally isolated chromosomal region that forms relatively few contacts overall, but those contacts it does make are biased toward other ribosomal RNA operons. Together, these patterns reinforce the hypothesis that ribosomal operons occupy spatially proximate positions within the nucleoid, likely reflecting their high transcriptional activity or coordinated regulation.

### Functionally Related Operons Cluster in Three-Dimensional Space

To minimize discretization artifacts arising from arbitrary bin boundary placement, we refitted the model across 32 coordinate shifts (16 forward, 16 backward, 3 kb per shift). This shift size approximates the average operon length, such that all shifts collectively span a full bin width in each direction. Interaction coefficients were re-estimated independently for each shift and then aggregated across shifts to identify pairs of genomic regions whose long-range interactions robustly influence overall bin behavior (see Methods for full details).

Aggregating across shifts improves resolution of biologically meaningful contacts: true long-range interactions persist across bin boundary changes, whereas interactions limited by binning artifacts may shift or weaken. We visualized these effects using “drop” and “rise” plots, which track changes in interaction coefficients resulting in a “rise” or “drop” of greater than 1 across different bin assignments. Network analysis of the aggregated interaction map (see Methods for details) revealed gene clusters that consistently interact topologically across shifts. While most bin-pair interactions occur at a basal level, a subset of long-range contacts remains significant and reproducible, highlighting non-random spatial clustering between specific distant genomic loci. The entire table of resulting coefficient rises and drops and associated genes is found in Supplemental Table 5, and resulting clusters and associated genes are found in Supplemental Table 6; we highlight a few key examples below.

Several prominent clusters correspond to operons or loci with coordinated physiological roles, including stress-responsive proteostasis and stringent response coupling, stationary-phase envelope remodeling, and σS-dependent metabolic and chaperone coordination. These clusters are visualized in Figure 3C–E, which shows both the overall interaction network and the subset of bin pairs with especially strong topological associations. To test for co-regulation among spatially clustered genes, we compared Spearman correlations of expression levels for gene pairs within these clusters against randomly chosen gene pairs.

The strongest example of functional clustering involves *mtlAD* (1,062,855 bp) and *ptsA* (2,579,636 bp), which show the highest within-cluster transcriptional correlation (ρ ≈ 0.16 vs. 0.002 background) and the largest coordinate-shift-induced drop in interaction coefficient (from 1.23 to −0.95; Figure 3D, green). MtlA is an Enzyme II complex mediating mannitol uptake and phosphorylation via upstream phosphoryl transfer from Enzyme I and HPr, while PtsA is a hybrid phosphotransfer protein containing HPr-, Enzyme I-, and IIA-like domains. PTS sugar utilization loci frequently evolve as modular units integrating substrate-specific transport with phosphotransfer and regulatory functions (Deutscher, 2008; Postma et al., 1993; Reizer et al., 1995). In this context, the spatial colocalization of *mtlAD* with *ptsA* may reflect functional coupling between sugar uptake and phosphorylation-state-dependent carbon signaling.

The second highlighted cluster links the *clpAS-serW* group (922,913 bp) with the *rpoZ-gmK-spoT* group (3,822,106 bp), which shows a strong rise in interaction coefficient (from −0.569 to 0.572; Supplementary Table 5; Figure 3D). We found a modest but significantly elevated transcriptional correlation relative to background (ρ ≈ 0.12 vs. 0.08 genome-wide). The functional composition of this cluster points to a shared role in coordinating proteostasis, transcriptional regulation, and stringent response signaling during stress, with (p)ppGpp as a plausible unifying signal: SpoT, RNAP (via RpoZ), and the ClpAP protease form a feedback loop activated under specific stress conditions (Gbolahan et al., 2025; Weichart et al., 2003, 2003). The combination of persistent three-dimensional contacts and modest expression correlation is consistent with a structural domain that is maintained constitutively but transcriptionally activated only upon stress.

A third cluster shows stronger transcriptional coupling (ρ ≈ 0.10 vs. 0.004 background; Figure 3F, pink), linking *yedACI–dgcQ* (2,029,539 bp) with *gadC–yddY–digH* (1,568,954 bp); the interaction coefficient for this cluster dropped from 1.52 to −0.059 upon coordinate shifting. These genes are associated with stationary phase and envelope stress, suggesting a shared role in coordinating acid resistance, membrane transport, cell division, and c-di-GMP signaling as cells exit exponential growth (Deutscher, 2008; Postma et al., 1993; Reizer et al., 1995). Spatial clustering may support localized c-di-GMP signaling through DgcQ, linking membrane-associated proteins and divisome components within the same chromosomal topological domain, or may reflect co-regulation by a common factor. The elevated expression correlation suggests potential co-induction during stress, though further experimental work would be needed to establish direct functional coupling.

Taken together, these clusters illustrate how spatial genome organization can group functionally related genes even when they are far apart along the chromosome. Mu-seq reveals these functional neighborhoods with sensitivity beyond what linear genome organization or expression correlation alone would predict, implicating chromosome architecture as an organizing principle of the bacterial stress response.

### Genomic Features Shaping Mu Insertions and Contacts

We next examined how Mu insertions and contacts relate to underlying chromatin features, using previously published hidden Markov model (HMM)-based chromatin states from Amemiya et al. 2022 (Amemiya et al., 2022b). The HMM was trained using IPOD-HR data and RNA pol ChIP-seq in wild-type cells and those lacking individual nucleoid-associated proteins. The six states in the resulting model integrate known transcription factor binding sites, promoter density, and NAP occupancy (Table 1, Figure 4). These states range from highly active (Active Regulatory) to closed EPOD-enriched regions (Closed 1 and 2), with intermediate states representing open or neutral chromatin (Open 1 and 2, respectively), and one state corresponding to a feature-poor, chromatin desert (Inactive).

**Figure 4.**
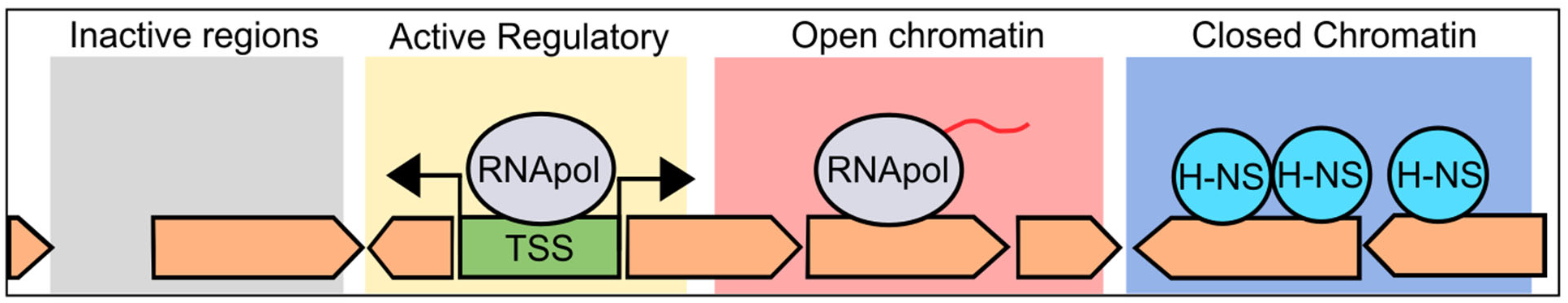
Chromatin state regulatory landscape. Schematic illustrating the mechanistic interpretation of six chromatin states, illustrating their relationship to transcriptional activity, accessibility, and protein occupancy, providing a framework for interpreting the contact patterns observed by Mu-seq.

**Table 1.** Chromatin State in *E. coli* characteristics.

| **HMM Class** | **Colloquial Name** | **Defining Features / Description** | **State Size (bp)** |
| --- | --- | --- | --- |
| 0 | Open 1 | Depleted for EPODs, transcription factors, and promoters. Low enrichment for H-NS and moderate Fis. | 1,124,160 |
| 1 | Open 2/Neutral | Slightly depleted for EPODs, promoters, H-NS, and TFs; near-average Fis levels. | 831,330 |
| 2 | Closed 1 | Strongly enriched for EPODs, TFs, and IHF, with exceptionally high H-NS occupancy (~3× genome average) and reduced Fis. Represents canonical H-NS–silenced heterochromatin. | 435,010 |
| 3 | Closed 2 | Strongest EPOD enrichment, strongly depleted for TFs, promoters and IHF; intermediate H-NS and Fis levels. Likely corresponds to repressed structural regions (e.g., long-range silencing domains). | 412,655 |
| 4 | Inactive | Depleted for nearly all features (EPODs, TFs, promoters, and DNA-binding proteins), despite being the largest state. | 1,326,280 |
| 5 | Active Regulatory | Strongly enriched for TFs, promoters, and IHF; moderate H-NS and highest Hfq/HU enrichment. Likely marks transcriptionally active and regulatory hubs. | 512,220 |

We constructed a genome-wide chromatin state map, binned to 10 kb, assigning each bin the identity of its most enriched state (Figure 5A). Unlike what we have observed in *Streptomyces venezuelae* (Ramirez et al., 2025) where closed states concentrate in chromosomal arms and open states in the core, no clear spatial segregation of states is apparent in *E. coli*, though Closed 1 appears visually enriched near the Ter region.

**Figure 5.**
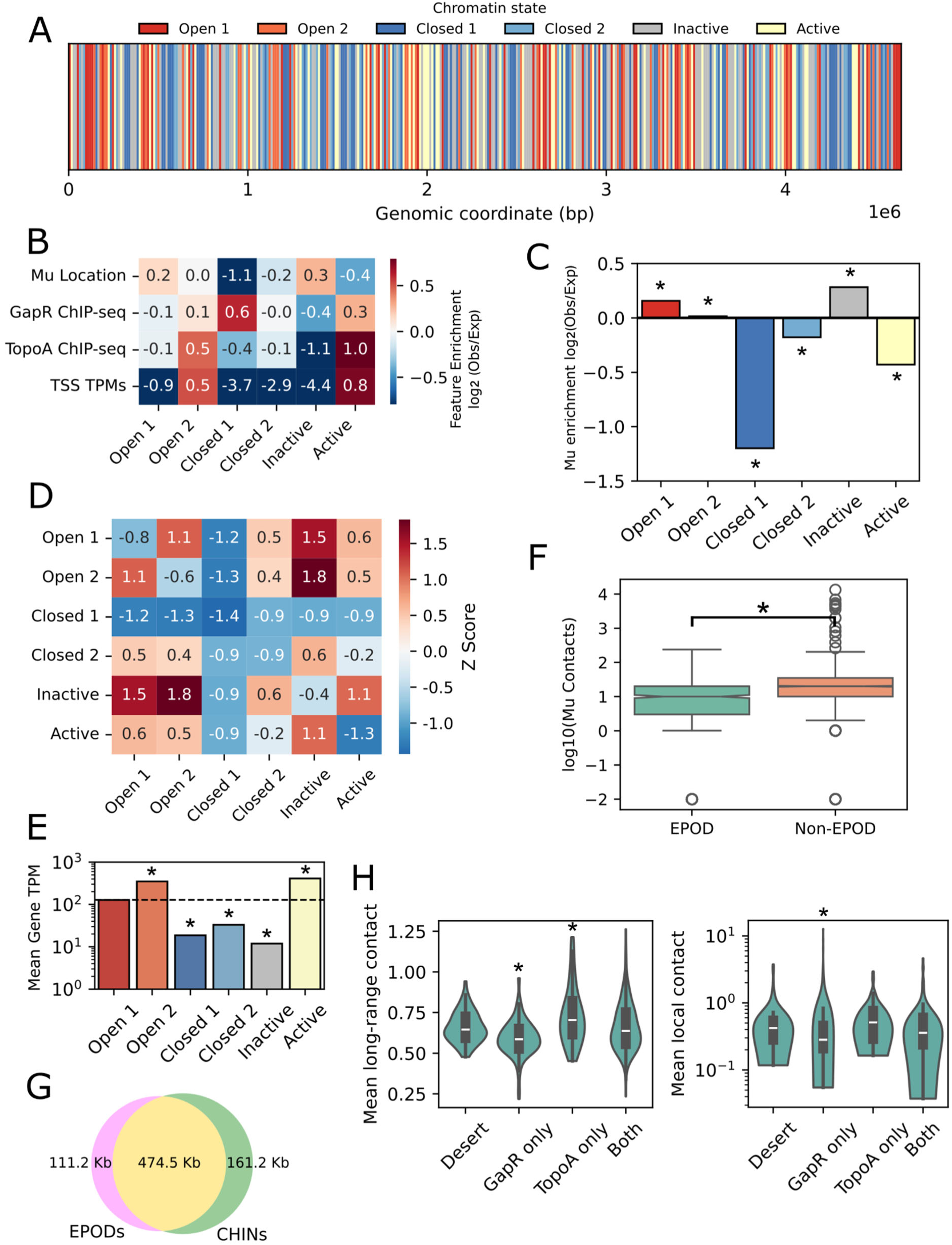
Structural features and protein occupancy explain variation in Mu transposition and contacts. **A)** Genome-wide map of HMM-based chromatin state annotations binned at 10 kb, showing the distribution and relative abundance of each chromatin state across the *E. coli* chromosome (for each bin, the most abundant state is shown). **B)** Summary matrix of enrichment values for genomic features-including Mu source and destination locations - across the six chromatin states. **C)** Barplot showing enrichment and depletion of Mu insertion locations in each chromatin state relative to expected frequencies based on genomic proportion. Asterisks indicate statistical significance. **D)** Z-score normalized matrix of non-local (≥92 kb) Mu contact frequencies between chromatin state pairs, adjusted for state size. **E)** Mean gene expression (TPM) by chromatin state. Asterisks indicate states with significantly different expression than Open 1 (Wilcoxon rank-sum test, FDR < 0.05). **F)** Box-and-whisker plot of the odds ratio for Mu insertion within EPODs relative to the rest of the genome. Notches indicate 95% confidence intervals around the median. **G)** Venn diagram showing the genomic overlap between EPODs and CHINs (Gavrilov et al., 2025) in *E. coli*. **H)** Violin plot comparing mean normalized Mu contact rates across bins classified by the presence of GapR binding sites, TopoI binding sites, both, or neither (desert). Asterisks indicate significance relative to the desert category.

To determine whether specific chromatin states are preferentially associated with Mu integration, we compared the observed frequencies of Mu locations in the uninduced and induced libraries against their expected genomic abundance, accounting for the size of each state. Enrichment and depletion were quantified as log₂(observed/expected) ratios and significance was tested using two-sided exact binomial tests, in which the null probability for each state was the fraction of its genomic abundance (Figure 5B–C). Mu insertions were significantly enriched in the Inactive state (log₂ enrichment = +0.29, p < 1×10⁻^10^) and modestly enriched in Open 1 (log₂ enrichment = +0.16, p < 1×10⁻^10^), while Closed 1 showed the strongest depletion (log₂ enrichment = −1.2, p < 1×10⁻^10^), followed by Active chromatin (−0.43, p < 1×10⁻^10^).

These patterns suggest that Mu transposition is shaped by chromatin structure, favoring accessible regions and disfavoring areas of dense protein occupancy, whether transcriptionally active or NAP-enriched.

We next assessed the enrichment of long-range interactions between chromatin state pairs, restricting the analysis to long-range interactions (≥92 kb apart, i.e., at least two bins). Contact counts between all state pairs were z-score transformed to normalize for state size, enabling direct comparison (Figure 5D). State pairs involving open or inactive chromatin (Open 1, Open 2, and Inactive) showed the strongest positive z-score deviations, indicating a propensity for long-range contacts, while Closed 1 was depleted for contacts both with itself and with other states, consistent with high protein occupancy restricting Mu transposition.

Mean gene expression levels further differentiated chromatin states, confirming Active as the most transcriptionally active (Figure 5B, E). Wilcoxon rank-sum tests indicated significantly higher median RNA transcript abundances (quantified in transcripts per million [TPMs]) in Active and Open 2 states relative to Open 1 (FDR < 0.05).

Notably, Open 2 combines high transcriptional output with nearly neutral Mu contact enrichment (score ≈ 0.01), whereas we observed substantial contact depletion in Closed 1 (enrichment score = –1.2) and Active (enrichment score = –0.43) states (Figure 5E). Together, these contrasts indicate that chromatin accessibility, rather than transcriptional activity per se, governs Mu insertion and contact propensity - consistent with prior ATAC-seq-based observations in *Caulobacter crescentus* (Melfi et al., 2021).

### Supercoiling, Chromatin State, and Mu Contacts

Building on the chromatin state analysis above, we cross-referenced Mu insertion sites with extended protein occupancy domains (EPODs) (Freddolino et al., 2021) and chromatin insulated regions (CHINs) (Gavrilov et al., 2025) to test the hypothesis that Mu transposition is strongly influenced by chromatin accessibility. Mu contacts were markedly depleted in protein-occupied regions (odds ratio = 0.22, p < 0.001), consistent with the insulated and transcriptionally silent nature of these domains (Figure 5F). This depletion further distinguishes structural accessibility from transcriptional activity: as noted above, Open 2 is associated with high gene expression yet is neither enriched nor depleted for Mu contact rates, confirming that Mu transposition reflects physical accessibility rather than simply transcriptional activity. Consistent with their molecular definitions, EPODs (characterized by dense protein occupancy) and CHINs (characterized by distinctive local DNA–DNA contact patterns) overlap extensively (63.5%) (Figure 5G), reinforcing the view that both mark transcriptionally silent, structurally insulated domains with similar molecular identities. Together, these findings demonstrate that Mu-seq reports on the structural and accessibility landscape of the chromosome independently of transcriptional state.

To investigate the relationship between local DNA supercoiling, chromatin state, and chromosome conformation, we integrated recently published GapR and Topoisomerase I (TopoA) ChIP-seq data (Fu et al., 2024) with *E. coli* chromatin states and Mu contacts. Analysis of ChIP-seq peaks across chromatin states revealed contrasting enrichment profiles for positive and negative supercoiling markers (Figure 5B). Topo I, which binds at sites of negative supercoiling, was highly enriched in the Active state and moderately enriched in Open 2, but depleted in Inactive and Closed 1 states, tracking closely with transcriptional activity. GapR, which binds at positive supercoils, showed the opposite pattern: moderate enrichment in Closed 1 and moderate depletion in Inactive chromatin. These findings indicate that negative supercoiling is maintained in transcriptionally active regions, while positive supercoiling accumulates preferentially in compact, closed chromatin domains (Royzenblat & Freddolino, 2024), consistent with findings reported in Fu et al., 2024 (Fu et al., 2024).

To explore how supercoiling-sensitive protein binding correlates with Mu contacts, we calculated the frequency of ChIP peak presence against Mu contacts per bin. GapR binding showed an inverse association with contact frequency (Pearson r = –0.188), with GapR peaks more often found in low-contact bins. Conversely, Topo I enrichment showed a positive correlation (Pearson r = 0.219), indicating that negatively supercoiled regions tend to occupy higher-contact bins. We then divided the chromosome into categories based on peak occupancy at 10 kb: bins bound by GapR-only, Topo I-only, both proteins, or neither (desert); and quantified Mu contacts normalized by observed/expected values at two scales: local (≤20 kb) and long-range (≥90 kb) (Figure 5H). GapR-only bins showed the highest mean local contact frequency (0.712) and the lowest mean long-range contact frequency (0.593), both significantly different from deserts (local FDR = 0.0071, long-range FDR = 0.00006). TopoA-only bins had the highest mean long-range contact frequency (0.723), which was marginally higher than deserts (0.659, FDR = 0.058), with no significant difference in local contacts (0.637, FDR = 0.23). Bins bound by both proteins exhibited mean values similar to desert bins, with no significant differences detected.

These patterns are unlikely to reflect a simple inability of Mu to transpose within closed chromatin. Because our analysis uses observed/expected normalized contact frequencies, it decouples contact propensity from raw chromatin accessibility and protein occupancy. Instead, the data support a model in which supercoiling state shapes the physical organization of the chromosome, with positive supercoiling favoring local compaction and negative supercoiling promoting long-range accessibility. Mu-seq thus characterizes chromosomal regions by their topological accessibility and interaction dynamics, beyond what insertion frequency alone would reveal.

### Transcriptional Activity and Chromosome Contact Frequency

To test directly for an association between transcript levels and DNA contacts, we compared normalized Mu contact intensity with RNA-seq expression levels across 10 kb genomic bins from cells grown at 37C in MOPS-RDM (Morgan et al., 2025) (Supplemental Table 8). To account for differences in DNA abundance across the genome, we normalized mean RNA abundance per bin by Mu contact counts as a proxy for DNA abundance, enabling comparison across bins with varying DNA content. Smoothed RNA abundance and contact density follow broadly similar patterns along the chromosome (Figure 6A).

**Figure 6.**
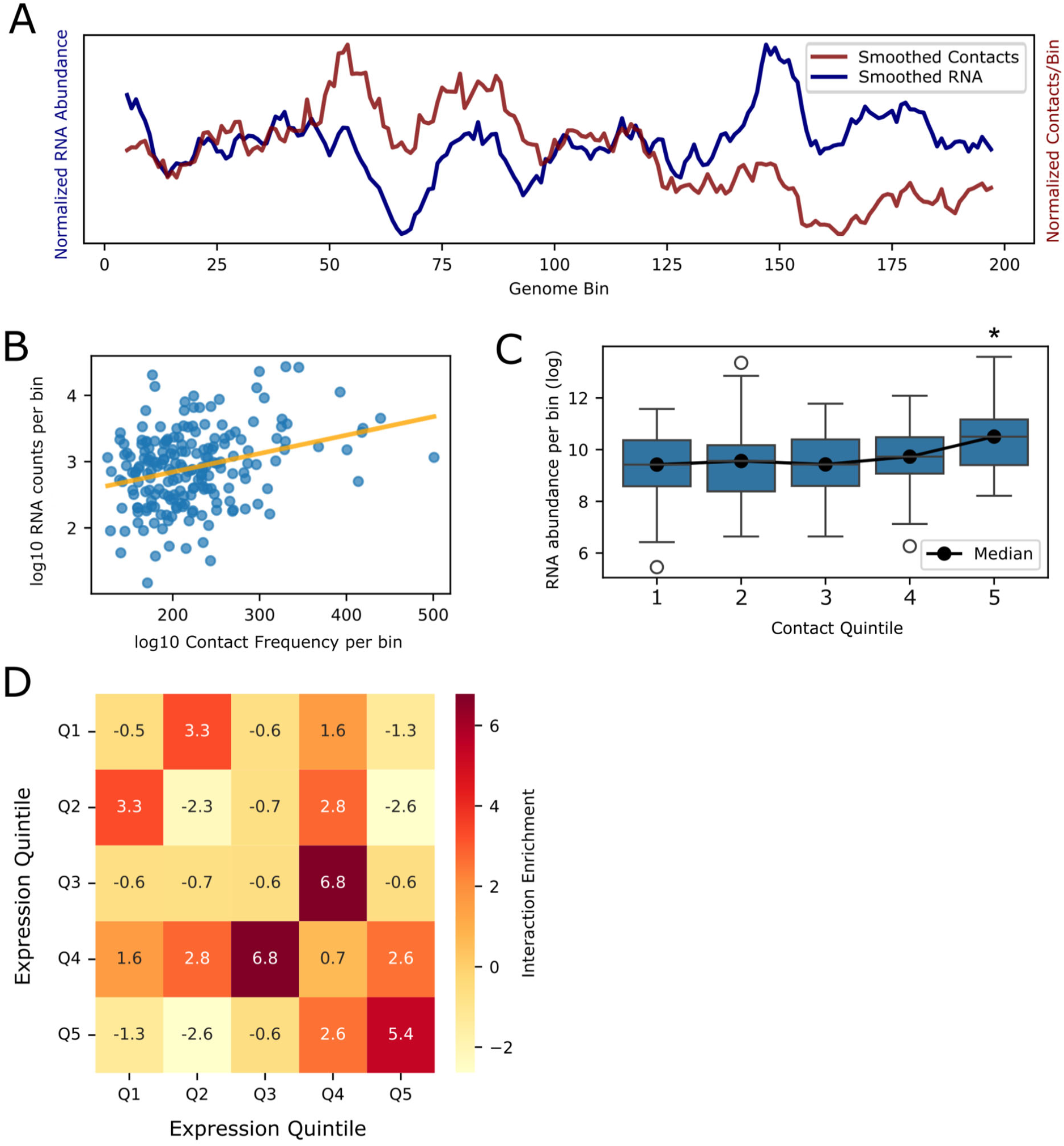
Relationship between chromosomal contacts and transcript levels. **A)** Smoothed genome-wide line plot comparing normalized RNA abundance to observed/expected-normalized Mu contact density per bin, showing that contact frequency correlates with RNA abundance. **B)** Scatter plot of log₁₀ RNA abundance versus log₁₀ total Mu contacts per bin, illustrating a positive relationship between transcriptional activity and contact frequency. **C)** RNA abundance stratified by contact quintile, showing enrichment in higher-contact bins. Asterisks indicate significance relative to quintile 3. **D)** Heatmap of mean non-local interaction coefficients between expression quintiles. Bins in the highest expression quintile preferentially interact with one another, while bins in the lowest quintile are depleted for non-local contacts. Enrichment is calculated as the mean interaction coefficient for each quintile pair divided by the global mean; values >1 indicate above-average interaction and values <1 indicate below-average interaction. Non-local interactions are defined as those >92 kb apart.

RNA abundance and contact counts were positively correlated (Pearson r = 0.32, p = 4.3e-6; Figure 6B). High-contact bins were enriched 2.7-fold for high-expression genes relative to random expectation, and Wilcoxon rank sum tests confirmed that RNA abundance was significantly higher within regions of high contact (p < 0.05). Stratifying contacts into quintiles revealed a consistent increase in RNA abundance in the highest-contact bins (Figure 6C).

We next asked whether bins involved in long-range contacts (≥92 kb) cluster by expression level - that is, whether genes with similar transcript abundances preferentially interact even when far apart on the linear genome. We assigned bins to expression quintiles based on the transcript abundance of contained genes, calculated the mean interaction coefficient for each quintile pair across all non-local bin pairs, and normalized each value by the global mean coefficient, producing a relative enrichment heatmap (Figure 6D).

Bins in the highest expression quintile (Q5) showed strong enrichment: Q5–Q5 contacts had mean interaction coefficients approximately 4.5 times the global average, indicating robust spatial clustering among highly expressed regions. Positive enrichment also extended to Q5–Q4 and Q5–Q3 contacts, suggesting that transcriptionally active regions preferentially co-localize in three-dimensional space across large genomic distances. Conversely, Q1–Q1 interactions were depleted relative to the global mean, and off-diagonal contacts involving Q1 bins—including Q1–Q5—also showed low or negative enrichment, supporting spatial segregation between active and inactive transcriptional domains. Bins in intermediate quintiles (Q2–Q4) showed moderate interactions with each other and with Q5, consistent with a gradation in spatial clustering along the expression spectrum.

Collectively, these results indicate that the spatial organization of the genome is influenced by transcriptional activity: highly transcribed regions form spatial clusters that span long linear distances, while low-expression regions remain comparatively isolated. This pattern is consistent with models of transcription factory formation and active/inactive compartments described in *E. coli* and other systems (Endesfelder et al., 2013; Ladouceur et al., 2020; Le et al., 2013).

## Discussion

Complementary experimental approaches have revealed different structural aspects of the *E. coli* chromosome: ribosomal RNA operon clustering is detected by fluorescence microscopy and Mu-based methods but not Hi-C, while macrodomain boundaries are clearly visible in recombination-based assays and Hi-C (Royzenblat & Freddolino, 2024). Mu-seq adds to this body of work through a fundamentally different physical principle – proximity detection via transposition in live cells rather than chemical crosslinking. A recurring theme across recent models is that the nucleoid must satisfy competing physical constraints: strong compaction yet high internal mobility. Differing observations regarding specific contact frequencies among Hi-C, microscopy, and Mu-based approaches likely reflect method-specific sensitivity to transient versus stabilized proximities, protein-occluded DNA, and population averaging. In this sense, different assays may capture different projections of a heterogeneous ensemble rather than a single fixed architecture.

Our results can be organized into three layers. First, we see the expected baseline behavior for a polymer: very high rates of short-range contacts that drop off with distance, consistent with basic polymer physics (Figure 2E–F; (Royzenblat & Freddolino, 2024)). A similar steep distance-dependent decay has also been observed in low-induction λ recombinase experiments, where recombination events are strongly enriched between nearby genomic sites (Valens et al., 2004); this reflects a general property of chromosomal polymer organization captured by recombination- and transposition-based proximity assays rather than a feature unique to Mu (it is perhaps notable that saturation of within-macrodomain hopping was observed in the λ based study noted above, but only at high levels of induction). Second, on top of that baseline, contacts are strongly shaped by chromatin state – specifically, how open or closed the DNA is, how much protein is bound, and how actively a region is being transcribed (Figures 5–6). Third, beyond both of those patterns, we find specific, long-range associations between particular loci and operons. For example, connections among *rrn* genes and clusters linked to stress and physiology (Figure 3C–F), suggesting that functional neighborhoods exist in the chromosome even when macrodomain-scale structure is weak.

The *rrn* cluster provides an important biological anchor for interpreting these functional neighborhoods. In prior work, long-range clustering of the dispersed ribosomal RNA operons was detected by fluorescence microscopy and independently recovered by Mu-seq, indicating that this network is not simply a statistical enrichment in an ensemble contact map. Moreover, *rrn* clustering is reversible and regulated: elevated sigma-H or FecI disrupts *rrn* operon clustering, whereas increased sigma-D suppresses this effect and restores clustering (Ho & Harshey, 2025). The anti-clustering activity of sigma-H requires interaction with core RNA polymerase and DNA binding but not sigma-H-dependent transcriptional activity, arguing that long-range *rrn* organization is not merely a passive consequence of high transcriptional output. Instead, the transcriptional apparatus can regulate chromosome topology independently of transcript production itself. Mu-seq now suggests that this principle may extend beyond ribosomal RNA operons to additional loci involved in stress adaptation, envelope regulation, and carbon utilization. We therefore propose that bacterial chromosome architecture contains dynamic functional neighborhoods whose assembly is coupled to physiological state rather than being fixed solely by linear genome position or polymer behavior.

Both major models of bacterial chromosome structure - the axial-core model and the solenoidal/toroidal DNA-phase model (Kuzminov, 2024) - predict that a compact chromosome need not be rigidly compartmentalized, and that long-range contacts can be common if the chromosome reconfigures frequently (across a cellular population and/or temporally over the cell cycle) and if active DNA is preferentially exposed at the nucleoid periphery. This helps explain why Mu-seq detects substantially more long-range interactions than crosslinking-based methods. Because Mu transposition requires both physical proximity and DNA accessibility, it preferentially records contacts involving exposed, dynamic DNA, whereas crosslinking methods may overcount frequent short-range and protein-stabilized interactions. Consistent with this, we observe faint CID-like boundaries but no macrodomain-scale plateaus (Figure 2A–D), and within-domain contacts are only modestly enriched relative to between-domain contacts (Figure 2C– D), arguing against a rigidly compartmentalized chromosome and pointing instead to a broadly interactive polymer with boundaries that are real but sharp, local, and transient.

Our chromatin state analyses reinforce this picture and reveal that “insulation” reflects both three-dimensional organization and DNA accessibility. Open and Inactive chromatin states show the most long-range contacts (Figure 5D), while Closed 1 regions, enriched for H-NS, are strongly depleted for both Mu insertions and contacts (Figure 5B–D). EPODs are similarly depleted for Mu signal (Figure 5F), consistent with the idea that densely protein-occupied DNA forms a mechanically and topologically constrained phase of the chromosome. Depletion in EPOD regions therefore likely reflects both reduced physical proximity *and* reduced accessibility for the transposition machinery: protein-rich compact domains act as barriers both to rearrangement and to topological access, while more open chromatin disproportionately contributes to long-range connectivity.

Transcription adds a further layer of organization. Highly transcribed bins preferentially contact each other over long genomic distances (Figure 6D), while the least transcribed bins show fewer long-range contacts, and RNA levels correlate positively with Mu contact frequency overall (Figure 6A–C). Within an axial-core/bottlebrush framework, this suggests that actively transcribed loci may be pushed toward the outside of the nucleoid, where DNA is more accessible, consistent with reports that RNA polymerase localizes to the nucleoid periphery and even the cell edge (Chai et al., 2014; Stracy et al., 2015). Kuzminov’s stratified nucleoid model (Kuzminov, 2024), which proposes a peripheral transcription zone, similarly predicts that highly expressed loci associate with each other at long range even when macrodomain boundaries are weak.

Our Mu data do detect stronger within-macrodomain contact correlations than between-macrodomain correlations (Figure 2D), as well as cohesion in the Ori region and relative isolation of Ter, indicating that macrodomain-scale features are present but subtle. Both patterns - macrodomain barriers prominent in Hi-C and broad long-range connectivity with expression-linked clustering prominent in Mu - likely reflect real features of a dynamic, heterogeneous chromosome population whose visibility depends on what each assay captures most efficiently.

Integrating our observations with prior literature, we speculate that rather than discrete insulated compartments, the *E. coli* chromosome could consist of overlapping structural and functional gradients: a dense axial scaffold enriched in protein-bound and low-expression regions, surrounded by dynamically extruded loops, with those loops further subdivided into inactive regions (which likely stay closer to the core) and active regions. Functionally, subsets of these peripheral or accessible loops can form long-range associations among loci with related physiological roles, creating neighborhoods that are not necessarily contiguous along the linear genome. A schematic representation of this hierarchical organization is shown in Figure 7, showing both larger-scale loops arising out of the axial core, and more frequent excursions even within each loop between more central (low expression, lower accessibility) and more peripheral (higher expression, higher accessibility) local environments. These conformations, furthermore, are simply representative of a large ensemble of possibilities present in any cellular population, with likely differences in contacts between different loops depending upon the stage of chromosomal replication/cell division and cell-to-cell variations within a large space of possible conformations even within each replication state. The most parsimonious explanation connecting our observations with those obtained recently using other methods is that crosslinking methods emphasize frequent, short-range, protein-stabilized contacts, while Mu requires proximity plus accessibility, enriching for transient, topologically exposed contacts that bridge distant loci.

**Figure 7.**
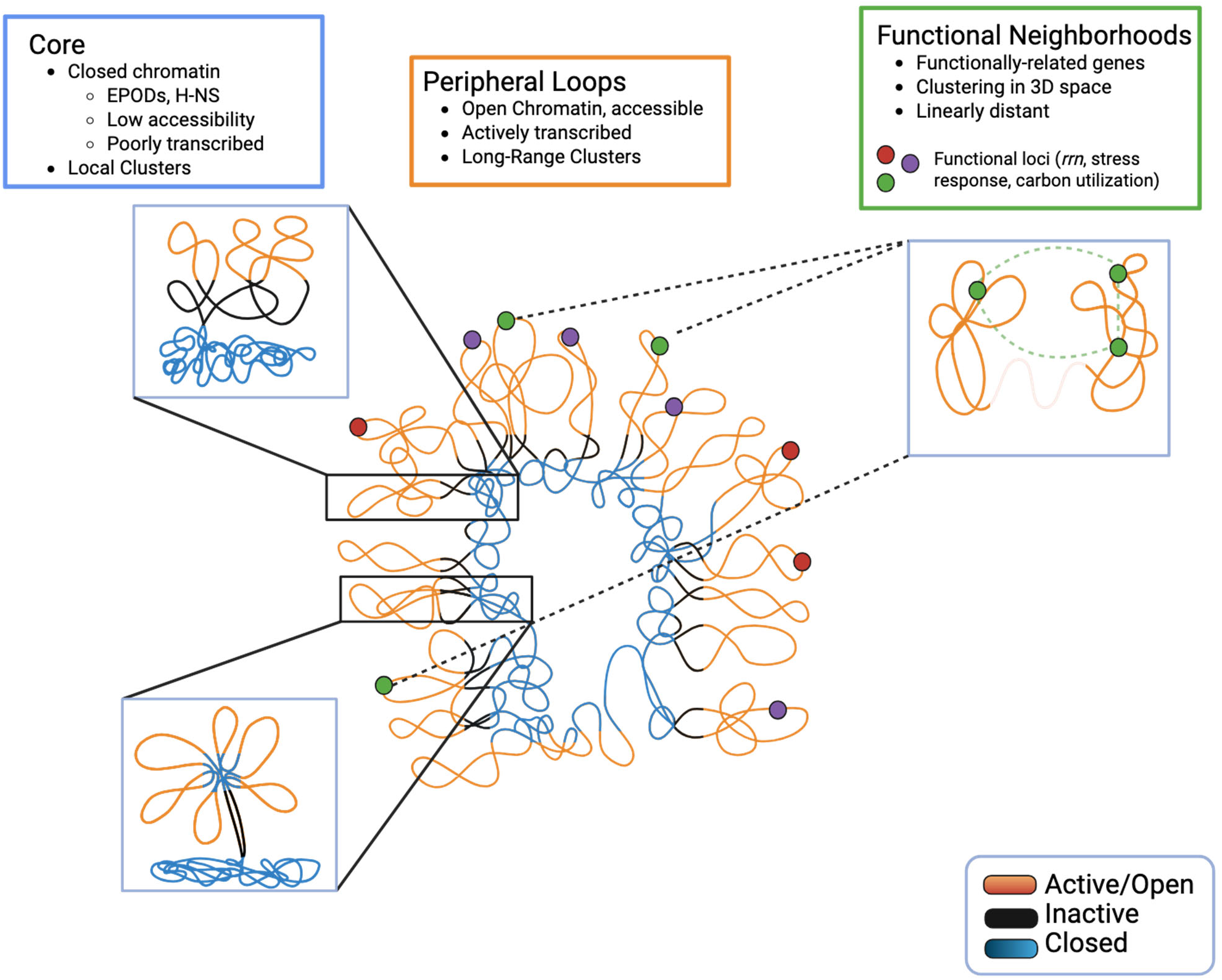
Schematic model of *E. coli* chromosome organization as revealed by Mu contact mapping and chromatin state analysis. Conceptual model of *E. coli* nucleoid architecture as inferred from Mu-seq contact mapping and chromatin state analysis. The dense core of the chromosome comprises closed chromatin regions – EPOD and H-NS enriched domains – characterized by local interactions and depletion of Mu insertions. Extending from this core are dynamically extruded loops formed by Active and Open chromatin regions, which carry higher transcriptional output and preferentially cluster in three-dimensional space, enabling long-range interactions among transcriptionally active loci. An Inactive chromatin zone separates these peripheral loops from the insulated interior. Functional neighborhoods represent long-range associations among loci with related physiological roles, such as stress response, carbon utilization, or ribosomal RNA production, that can occur despite large separation along the linear genome. These neighborhoods may arise through transient or regulated contacts between accessible chromosomal loops rather than through fixed compartment boundaries. This architecture agrees with an axial core model of nucleoid organization (Mäkelä & Sherratt, 2020) in which *E. coli* chromosome structure contains a series of loops extruded from a MukBEF axial core; furthermore, comparing the length scales of variations in accessibility to the necessary loop sizes indicates that within a single topological loop out of the MukBEF core, multiple excursions between active and inactive states occur (e.g., bottom left). Our findings are also compatible with microscopy-based models in which H-NS-bound DNA forms centrally localized clusters (Wang et al., 2011), while active transcriptional regions extend toward RNAP- and ribosome-rich peripheral zones that may facilitate transcript production, diffusion, and translation (Bakshi et al., 2012, 2015; Endesfelder et al., 2013).

Our model is consistent with microscopy-based evidence for radial organization of the *E. coli* nucleoid. In this view, H-NS-bound and other protein-occupied regions contribute to a dense internal scaffold, whereas accessible and transcriptionally active DNA is preferentially exposed on dynamic peripheral loops. Prior imaging showed that H-NS localizes to discrete clusters near the cell center, unlike many other nucleoid-associated proteins, and that H-NS can mediate long-range DNA interactions (Wang et al., 2011). Our observation that H-NS-enriched, EPOD-like, and other protein-occupied domains are depleted for Mu insertions and contacts provides an orthogonal contact-mapping signature of this centrally compacted, relatively inaccessible chromatin fraction. Conversely, the enrichment of long-range Mu contacts among open and transcriptionally active regions suggests that active loci are more topologically exposed and more likely to encounter one another in the peripheral nucleoid environment.

This framework also generates concrete, testable predictions. If highly expressed genes cluster through a peripheral transcription zone, acute transcription inhibition with rifampicin should selectively weaken long-range contacts between highly transcribed genes (e.g., gene pairs in the upper quintile of expression levels) while leaving the basic polymer distance law intact. If EPOD/CHIN/H-NS regions form a physically closed phase, disrupting H-NS or EPOD formation should increase Mu accessibility in closed chromatin and shift the balance of local versus long-range contacts. Testing these predictions would help determine whether the stratification seen by Mu primarily reflects spatial geometry (core-periphery organization), topological state (supercoiling and loop constraints), or some combination.

### Technical Limitations and Future Directions

Implementing Mu-seq as a genome-wide all-to-all contact probe revealed an inherent information bottleneck that points toward a clear path for improvement. Although we detected 198,294 source and 234,059 destination barcode locations - providing theoretical ∼23 bp resolution - only a small fraction could be assigned as matched source-destination pairs under stringent filtering. This sparsity arises primarily from library complexity outpacing sequencing depth: with ∼100,000 unique barcodes each present approximately 72,000 times in the population, a single round of transposition generates ∼7.2 billion unique integrants. At 25× coverage, the probability of recovering any given barcode from source to destination is only ∼16% under ideal conditions; our actual match rate was ∼4.5%, further reduced by splitting sequencing depth across seven samples.

Crucially, this bottleneck is technical rather than fundamental, and its solution is straightforward: reducing library complexity at the input stage. Targeting ∼10,000– 20,000 barcodes rather than ∼100,000 - while maintaining low population density and our current sequencing depth of 1.2 billion reads - would increase barcode recovery probability roughly fivefold, bringing matched source-destination pair recovery well above 50% under realistic conditions. This single change would transform Mu-seq from a resource that detects global organizational principles into one capable of high-resolution, locus-specific contact mapping across the full genome.

The present dataset already demonstrates the biological power of the approach despite these constraints: we detect CID-like frontiers, preferential contacts among open chromatin states, depletion in EPODs and H-NS-bound regions, and long-range clustering of functionally related operons - features not accessible to crosslinking-based methods. The results reported here thus represent a lower bound on what Mu-seq can achieve, with a well-defined and experimentally tractable route to substantially higher resolution and sensitivity.

## Supplementary Material

**Supplemental Figure 1.**
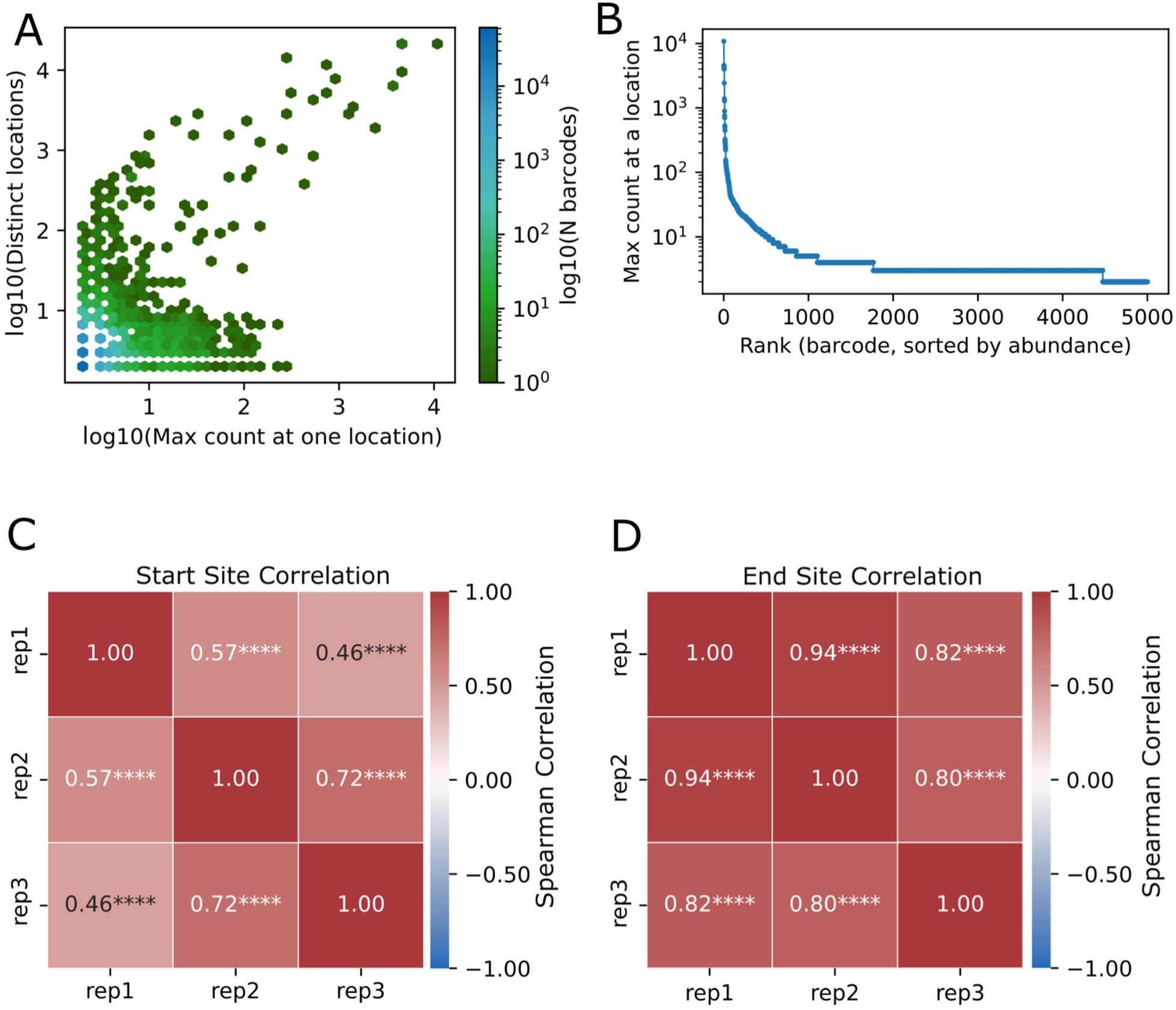
Barcode distribution and diversity metrics across the Mu transposition libraries. **A)** Range and abundance of each barcode plotting how often each barcode occurs in one location versus distinct locations, highlighting diversity and indicating the presence of rare hyperactive barcodes. **B)** Rank abundance demonstrates the non-redundant nature of the barcode libraries. **C-D)** Heatmap values indicate Spearman correlations between binned Mu insertion profiles across replicates. C shows correlation between start sites, D between end sites. Significance stars indicate nominal correlation p-values: *p < 0.05, **p < 0.005, ***p < 0.0005, ****p < 0.00005.

**Supplemental Figure 2.**
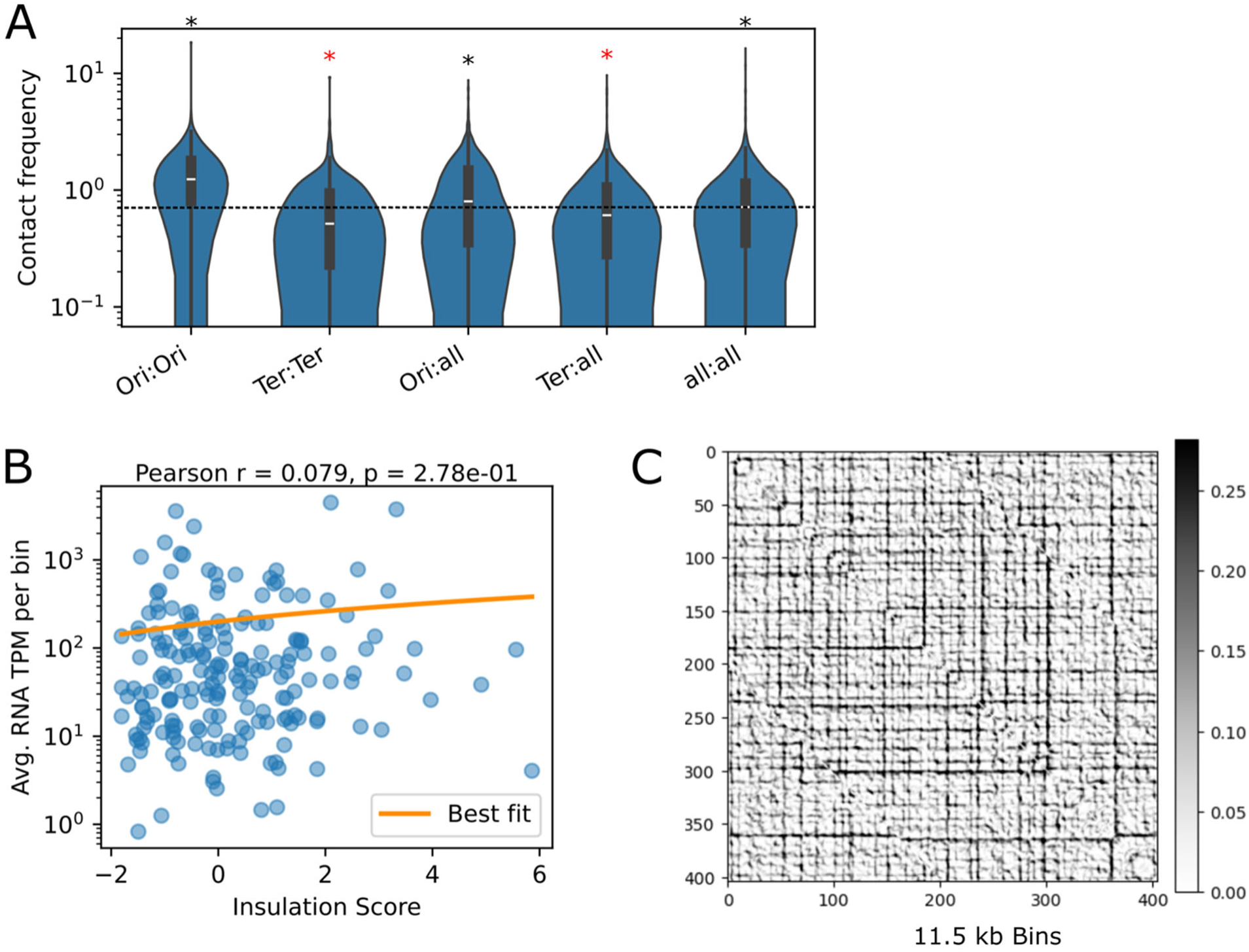
Additional analysis of domain boundaries and transcript levels. **A)** Violin plots show the distribution of observed/expected-normalized Mu contact frequencies among Ori:Ori, Ori:All, Ter:Ter, Ter:All, and All:All bin pairs. Dashed line represents the median contact frequences for All:All bin pairs, asterisks indicate significance, with red star denoting the contact pairs which fall below All:All and black asterisks denoting those that fall above. P values in the accompanying table were calculated using two-sided Wilcoxon rank-sum test comparing each category against the all:all background, indicating enrichment of Ori-associated contacts and depletion of ter-associated contacts (Supplemental Table 2). **B)** Scatter plot of RNA abundance for each 23 kb genomic bin as a function of its insulation score from Figure 2B. The fitted regression line and correlation statistics are shown, indicating a weak and non-significant association between insulation score and RNA abundance across the bins. **C)** Frontier index analysis shows, at an 11.5 kb resolution, the location of insulating boundaries and strong transitions in chromosome contact density. The frontier signal highlights the regions where there is a large change in the contact patterns between adjacent bins.

### Supplementary Data/Table Captions

**Supplemental Table 1. Tabulation of all contacts**. Post-processing identification of all detected contacts from Mu-seq, with replicates merged. Source column contains Mu start coordinates, destination column contains Mu end coordinates, count denotes the number of occurrences (reads) where these coordinate interactions were detected.

**Supplemental Table 2. Contact_corr_obs_vs_exp.** Observed/expected-normalized Mu contact frequencies were calculated among Ori:Ori, Ori:All, Ter:Ter, Ter:All, and All:All bin pairs. Total contacts per category are provided in the table, alongside P values calculated using two-sided Wilcoxon rank-sum test, plus U test values additionally included.

**Supplemental Table 3**. **Short-range enrichment of Mu hops.** Permutation test comparing short-range Mu hop enrichment to long-range.

**Supplemental Table 4. Distance_bin_enrichment.** Observed/expected Mu hop enrichment, as well as short- and long-range contacts.

**Supplemental Table 5. All shifts with genes.** Coefficient rises and drops across shifted bin assignments. Bin_label column denotes the interacting bins. From_shift and to_shift indicate the starting and ending coordinate shift distance relative to the genes starting coordinates. From_coef and to_coef refer to the interaction coefficient calculated at the “from_shift” location and in the to_shift location, respectively. Gene_bin refers to the bin each gene is from, this corresponds to the bin_label IDs. Driver_gene refers to the identified gene driving the change in the interaction coefficient change. Event_direction distinguishes between shifts that yielded a rise in interaction coefficient or a drop in interaction coefficient, greater than 1.

**Supplemental Table 6. All Clusters.** Gene clusters from shift-aggregated interaction networks. All identified cluster IDs, genes occurring in the cluster, and gene start coordinates are found in respective columns.

**Supplemental Table 7. Primer summary.** Full sequences of primers used in the methods as described.

**Supplemental Table 8. Summary of datasets** used in this work. The “Dataset” column refers to the specific dataset referenced, the “Reference” column refers to the material reference used in the paper, and “DOI/Source” refers to the accession link for datasets.

**Supplementary Data 1**: Normalized correlations of transcript levels across conditions for all genes used in our analysis, derived from the COLOMBOS database (see Methods for details).

### Methods

#### Barcoded Mu Library Generation

Our barcoded Mu library was generated as exactly as in Ho and Royzenblat et al. 2024 (Ho et al., 2024), (see Figure 2).

#### Genomic DNA preparation

Genomic DNA from each sample was extracted following the protocol in (Ho et al., 2024). Following purification, samples were quantified using the Promega QuantiFluor® dsDNA System, a fluorescent quantitation assay. To enrich Mu-containing genomic DNA, a transposon footprinting based method was used.

#### U-Linker-based DNA Footprinting

##### U-Linker Synthesis

A 10x annealing buffer was prepared by combining 20 µL of 5 M NaCl, 10 µL of 1 M Tris-HCl (pH 8.0), 2 µL of 0.5 M EDTA (pH 8.0), and 68 µL of milliQ water.

The U-Linker was synthesized using primer P4614 as in Supplemental Table 1. NEB Cutsmart Buffer was used to anneal the Ulinker complementary ends. To anneal, 10 µL of 100 µM U-linker oligonucleotide was mixed with 5 µL of 10x Cutsmart buffer and 35 µL of water in a PCR tube, for a final volume of 50 µL, heated to 90°C for 2 minutes, and then gradually cooled to 30°C at a rate of ∼2°C per minute. The tubes were snap-cooled on ice. The annealed oligo was then diluted to 100 µL final volume by adding 10 µL Cutsmart buffer and 35 µL milliQ water, then briefly vortexed. Subsequently, 5 µL of MspI enzyme was added and the mixture incubated at 37°C for 2 hours, followed by purification using Zymo Oligonucleotide Clean and Concentrate clean-up kit, eluting in 21 µL of buffer. For further annealing, 2 µL of 10x annealing buffer was mixed with 18 µL of the digested U-linker, heated to 80°C for 2 minutes, then cooled to 23°C at a rate of 0.1°C/sec, and maintained on an ice block until use.

##### Sample Digestion

Sample genomic DNA was adjusted to 1 µg per reaction, then digested in a 50 µL reaction containing 5 µL10x Cutsmart buffer, variable volume DNA (to reach 1 µg), water to 42 µL, and 3 µL Msp1, assembled on ice in PCR tubes. Reaction was incubated for 3 hours at 37 C in a thermocycler and held at 4 C until ready to use. A four base pair cutter was chosen that cuts outside of our right Mu end containing the barcode. DNA was purified using Zymo Clean and Concentrate −5 kit, and eluted in 10 µL.

##### Ligation of U-Linker to DNA Fragments

For ligation, on ice and in PCR tubes, 2.5 µL of freshly cut U-linker was mixed with 9 µL of purified digested DNA and 12.5 µL of 2x Quick Ligation buffer and mixed well; 1 µL of Quick Ligase (NEB M2200) was added, and reactions incubated at room temperature for 10 minutes. Ligation was terminated with 1 µL of 0.5 M EDTA.

##### Axygen Bead Purification

Ligation products were purified by adding 23 µL of thoroughly resuspended AxyPrep MAG DNA beads, followed by 5 minutes of incubation at room temperature. Beads and sample mix were washed twice with 200 µL of freshly prepared 80% ethanol, air-dried on a magnetic stand for 5 minutes, and DNA eluted with 23 µL TEe buffer. 20 µL of eluate was transferred to new tubes.

##### PCR Amplification for Illumina Library Preparation

PCR was performed to add Illumina-compatible sequences. Per sample, a reaction mix containing 12.5 µL water, 10 µL 5X NEB Q5 Reaction Buffer, 2.5 µL of P4537 (this is the i5 Mu primer, 10 µM) and P4599 (this is the i7 primer and binds to the U-Linker, 10 µM), 1 µL dNTPs, 0.5 µL NEB hot-start Q5 Polymerase, and 1µL USER enzyme, was mixed in PCR tubes, on ice. 20 µL of bead-purified DNA was added to each tube, for a total of 50 µL. Thermocycler conditions were: USER treatment at 37°C for 30 min, 95°C for 1 min, 4 cycles of (95°C for 15 s, 63°C for 15 s, and 72°C for 1 min); end with 72°C for 2 min and held at 4°C.

##### DNA Fragment Clean Up, Quantification, and Sequencing

PCR products were purified by adding 36 µL of thoroughly resuspended AxyPrep MAG DNA beads, followed by 5 minutes of incubation at room temperature. Beads and sample mix were washed twice with 200 µL of freshly prepared 80% ethanol, air-dried on a magnetic stand for 5 minutes, and DNA eluted with 23 µL 0.1X TE buffer. Libraries were then again prepped for PCR as above, this time using 10 µL premix NEBNext® Multiplex Oligos for Illumina® NEB #E6440S/L for primers, 5x Q5 buffer, 10 mM dNTPs, 20 µL cleaned library, Q5 polymerase, bringing the total volume to 50 µL. Libraries were purified as above with AxyPrep MAG DNA beads, this time using 0.5X total volume. With a final elution in 1x TEe buffer (10mM Tris (pH 8 or 7.5) and 0.1mM EDTA).

Size distribution of PCR products was visualized by electrophoresis on a 1% agarose gel, and average fragment size recorded. DNA concentration of each library was determined using the Promega QuantiFluor® dsDNA System. Libraries were further standardized using Agilent TapeStation Model 4150. Final libraries were run on an Illumina Novaseq X plus Instrument, paired-end 150 bp sequencing according to manufacturer’s protocols, ensuring sufficient read length to contain both the barcode at one end and genomic DNA at the other.

#### Bioinformatic Processing of Sequencing Data

Analyses were performed using custom Python and R scripts. Software versions and package dependencies are provided in the accompanying conda environment (.yml). Several matrix-processing functions (SCN/ICE normalization, distance-law estimation, domainogram/directionality computations, and related utilities) were taken from the Koszul lab *E. coli* analysis codebase (https://github.com/koszullab/E_coli_analysis), originally developed for (Lioy et al., 2018). External datasets for cross-referencing Mu features were selected from experiments performed under comparable growth conditions, to match the conditions used for Mu library construction. A complete table can be found in Supplemental Table 2.

##### Read Alignment and Mapping of Insertion Sites

Using Cutadapt, barcode sequences were extracted from the Read 1 fastq files by identifying reads containing the Mu motif flanking the barcode (sequence), and extracting the 14 nucleotide N-region, corresponding to the unique barcode for each read. In footprinting experiments, the U-linkers incorporated an 8 bp random unique molecular identifier (UMI), detected in Read 2.For identification of where the Mu transposon is located in the genome, Read 2 files were filtered to retain only reads containing the U-Linker adapter sequence, which is expected immediately before the genomic DNA, which was trimmed as a 5′ adapter. At the 3’ end of the read, we may read into the right element of the Mu motif, and so that was included as a potential adapter, but not required. The retained sequences represented the genomic DNA adjacent to the Mu insertion. In footprinting experiments, the U-linkers incorporated an 8 bp random unique molecular identifier (UMI), detected in Read 2, to account for PCR duplicates. Following 5′-end trimming, trailing construct sequences were removed to recover UMI sequences from the remainder of each read. Information from UMIs, barcodes, and insertion coordinates was then matched for each read, retaining only those entries in which all three elements were present.

Paired-end reads were matched such that only read pairs present in both Read 1 and Read 2 files were retained for downstream analyses. Genomic DNA sequences from filtered Read 2 files were aligned to the appropriate reference genome (using bowtie2) to precisely identify integration sites; the genomic location nearest the start of each trimmed read was recorded as the Mu insertion site. For alignment, a composite reference was constructed consisting of the *E. coli* K-12 MG1655 reference genome (GenBank: U00096.3) and the modified Mu phage reference genome (see supp data). Because fragment length after restriction digestion is uncertain, insertion regions sharing a barcode were aggregated to within 200 bp, corresponding to the 70th percentile of Msp1 fragments. The complexity and diversity of libraries were assessed in three independent biological replicates and two technical replicates, with an initial bottleneck of ∼100K barcodes (determined by PFU).

The resulting information containing barcodes, UMIs, and aligned genome positions were deduplicated such that only one unique combination of each barcode, UMI, and coordinate was retained. To ensure unambiguous mapping, barcodes found at multiple insertion sites in the uninduced library were excluded.

In total, after pre-processing, **119,106 barcodes** were mapped to the *E. coli* genome covering **234,059 unique locations** and these were retained for subsequent analysis. These filtered data were used to map genomic contacts across the chromosome.

Data from all libraries, including both pre- and post-induction samples across biological replicates, were analyzed as follows. The genome was divided into bins based on the chosen resolution, producing bin identities for each read. Barcodes mapping to more than one location in the primary library were excluded to prevent ambiguity. The outcome was a raw transposition frequency matrix, constructed by counting the number of reads whose barcode (source bin) mapped to each possible destination bin in the secondary library sample, forming a contact matrix. Normalization and downstream significance testing were performed using custom or published methods tailored to the specific goals of the analysis.

##### Mu insertion enrichment

Normalized Mu insertion enrichment and depletion across the E. coli genome were calculated by dividing observed Mu insertion counts by per-bin barcode coverage at 1 kb resolution. Z-scores and quantile analysis identified bins with statistically significant enrichment or depletion, which were superimposed on genome coordinates annotated for Ori and Ter macrodomains; Mann–Whitney U tests assessed regional biases.

##### Opportunity-normalized transposition distance

Source–destination distances were computed for all transposition events using the shorter of the linear and circular distances. Distances were binned into five logarithmically spaced intervals and normalized by opportunity (the bp span of each interval) to account for the greater number of possible locus pairs at longer separations. To assess whether short-range contacts were enriched beyond the expected distance decay, observed opportunity-normalized rates were divided by the empirical decay expectation, and the mean enrichment difference between short-range and long-range bins was evaluated using a two-sided permutation test (10,000 permutations).

##### Contact Matrix Construction and Normalization

Mu transposition contact counts were aggregated into genome-wide matrices at 46 kb resolution. Raw count matrices were normalized using sequential component normalization (SCN) and rescaled to preserve total contact counts. For analyses requiring observed/expected (O/E) normalization (e.g., RNA–contact comparisons; Fig. 6), expected contact frequencies were computed from row and column marginals as

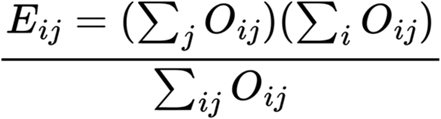

and O/E ratios were log-transformed, symmetrized by averaging reciprocal entries, with the diagonal set to zero.

##### Distance-Law Correction, Correlation Analysis, and macrodomain comparison

A genome-wide distance law *P(s)* was empirically estimated from the SCN-normalized matrix to model expected contact decay as a function of genomic separation. Each entry in the contact matrix was divided by the corresponding distance-law expected value to produce a distance-law-normalized matrix. Pearson correlation coefficients were then computed between the distance-law-normalized interaction profiles of all bin pairs, yielding a correlation matrix that captures similarities in genome-wide contact behavior (“guilt by association”). A second-order correlation matrix (correlation of correlation profiles) was additionally computed for visualization (Fig. 2C).

Macrodomain assignments were applied to Figure 2C matrix bins following published definitions (Valens et al., 2004). Pairwise correlation coefficients from the distance-law-normalized correlation matrix were stratified as: (i) canonical within-domain (same macrodomain) and (ii) between-domain (different macrodomains). Differences in mean correlation between within- and between-domain groups were assessed using two-sided permutation tests (10,000 label permutations) and two-sided Mann–Whitney U tests.

##### Directionality Index and Frontier Analysis

Directionality index (DI) profiles and frontier signals were computed using methods from Lioy et al, with the SCN normalized matrix binned to 10 kb as the input. The DI track at scale 40 (corresponding to ∼400 kb) was used for visualization. For the scale-dependent local contact signal, each genomic bin was assigned the sum of SCN-normalized contacts within a circular window of radius *n* bins, with *n* varied across scales. This produced a two-dimensional scalogram in which genomic position is shown on the x-axis, window size on the y-axis, and color indicates the relative summed local contact signal.

##### Distance Decay and Opportunity Normalization

Contact frequencies were quantified as a function of genomic distance using the empirically estimated distance law. To enable comparison across genomic separations, observed contact counts at each distance were normalized by “opportunity”, ie. the number of bin pairs separated by that distance, accounting for the circular chromosome topology.

##### Poisson Regression Modeling of Chromosomal Interactions

To systematically model interaction frequencies while controlling for confounding factors, binned contact counts (46 kb bins) were fit using a Poisson generalized linear model with LASSO regularization implemented in the R package glmnet. The model design matrix included: indicator variables for interaction type (one-sample versus paired two-sample), replicate identity and replicate-by-type interactions, short-distance indicators (bin distance = 1 or 2), per-bin effects (encoded separately for one-sample and paired data to reflect the different information content of each), and bin-pair identity terms to capture specific pairwise interaction strengths. Log-transformed per-bin DNA abundance (mean normalized coverage across replicates) and log-transformed inverse square-root genomic distance were included as offsets. The regularization path was evaluated over 251 lambda values (log-uniformly spaced from 1,000 to 0.01, plus lambda = 0), and the optimal penalty was selected via cross-validation. Fitted interaction coefficients at the bin-pair level were interpreted as the residual interaction strength after controlling for distance, DNA abundance, and technical covariates.

#### Methods for the gene-gene expression correlation matrix

The *E. coli* transcript-level expression data was downloaded as a compendium from COLOMBOS v3.0 (Moretto et al., 2016). COLOMBOS manually curates publicly available microarray data and homogenizes the experimental annotations. A “contrast” in COLOMBOS is defined as a paired test-versus-reference comparison of samples in the experiment context. Within a contrast, the expression score of each gene was calculated as the log_2_-fold-change of the test and the reference expression values.

To normalize expression changes for comparability, the modified z-scores were calculated for each contrast across all genes using the raw values from the COLOMBOS matrix. The modified z-score is defined as the difference between a gene’s raw expression value and the median of all values in the contrast, divided by the median absolute deviation of the contrast multiplied by a constant for asymptotically normal consistency. See the equation below for the modified z-score (*z*) calculation, where *X* is the raw COLOMBOS expression values for each gene (*i*) given a contrast, and *X̅* is the median of *X*.

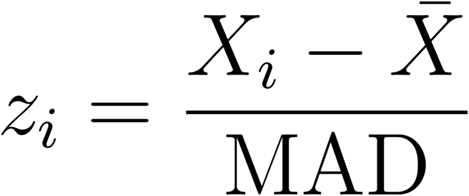

Where MAD = 1.4826 × *median*(|*X_i_* − *X̅* |)

To assess gene co-expression patterns, pairwise Spearman correlations were calculated for each pair of genes from the normalized gene-contrast matrix of modified z-scores using complete pairwise observations. All calculations for the gene-gene expression correlation matrix were performed in R.

##### Bin-Shift Analysis, Gene Network Construction, and Co-expression analysis

To identify long-range interactions robust to arbitrary bin boundary placement, the Poisson regression model was refit across 32 coordinate shifts (16 forward, 16 backward, each offset by ∼3 kb, approximately one average operon length). For each shift, interaction coefficients were independently re-estimated. Changes in bin-pair coefficients exceeding a magnitude of 1 were tracked across shifts, and the genes gained or lost from each bin at each shift were recorded to identify genomic features driving specific interactions.

Gene co-occurrence networks were constructed by connecting genes that appeared together across bin-shift events, with edge weights proportional to the number of co-occurrences. Networks were analyzed using standard graph metrics (degree, betweenness centrality) and community detection via greedy modularity maximization (NetworkX (Hagberg et al., 2008)). Network visualizations were restricted to the largest connected component where indicated.

To test whether spatially clustered gene groups from our network analysis exhibit correlated transcription, pairwise Spearman correlation coefficients were computed for genes within each identified cluster using a pre-computed genome-wide gene-gene expression correlation matrix (see above). Only gene pairs separated by ≥50 kb on the linear genome were included to exclude trivially co-regulated neighbors. Within-cluster median Spearman correlations were compared to the median correlation of each cluster gene with all genes outside the cluster, providing a background expectation.

##### Chromatin State Enrichment

Six-state HMM chromatin annotations were obtained from (Amemiya et al., 2022a). For each chromatin state, the expected frequency of Mu insertions was defined by the state’s genomic fraction (proportion of total base pairs). Enrichment was reported as the average log₂(observed/expected) across source and destination, and the average was computed. Global deviation from the null expectation was assessed by chi-square goodness-of-fit tests, and per-state significance was evaluated using two-sided binomial tests with Benjamini–Hochberg FDR correction.

##### Enrichment of features within chromatin states

Non-local Mu contacts (≥92 kb separation, corresponding to ≥2 bins at 46 kb resolution) were tabulated between all pairs of chromatin states. Contact counts were converted to Z-scores after normalizing for the genomic size of each state, enabling comparison of interaction propensities across states of different abundances (Figure 5D).

#### EPOD and CHIN Overlap and Mu Contact Depletion

EPOD annotations were obtained from (Freddolino et al., 2021) and CHIN annotations from (Gavrilov et al., 2025). Base-pair-level overlap between EPODs and CHINs was computed using interval tree logic, and the significance of the observed overlap was assessed by a randomization test (1,000 iterations) in which CHIN intervals were placed at random genomic positions while preserving their lengths. To quantify Mu contact depletion in protein-occupied domains, per-bin Mu contact values were classified as overlapping or non-overlapping with EPOD intervals (using bedtools intersect), and the difference in mean contact intensity was assessed using a circular-shift permutation test (1,000 random offsets) preserving spatial autocorrelation, yielding an empirical two-sided p-value.

#### GapR and TopoI ChIP-seq Integration

GapR and TopoI (TopoA) ChIP-seq peak data were obtained from Fu et al. (2024)(Fu et al., 2024). Peak counts were intersected with the six-state chromatin annotation, and enrichment was computed as log₂ (fraction of peaks in state / fraction of genome in state). To relate supercoiling markers to Mu contact patterns, 10 kb bins were categorized by peak occupancy (GapR-only, TopoI-only, both, or neither/“desert”), and mean observed/expected Mu contact frequencies were compared at local (≤20 kb) and long-range (≥90 kb) scales. Pairwise differences relative to desert bins were assessed using Mann–Whitney U tests with FDR correction.

##### Transcriptional Activity and Chromosome Contact Frequency

Bulk RNA-seq expression data were obtained from Morgan et al. 2025 (Morgan et al., 2025). Gene-level transcript abundances were mapped to genomic bins using strand-aware transcription start site (TSS) assignments. Per-bin RNA abundance was compared to per-bin Mu contact density using Pearson and Spearman correlations and visualized with rolling-window smoothed genome-wide tracks and scatter plots. RNA abundance was additionally normalized by per-bin DNA abundance (Mu contact counts as a proxy) to control for copy-number variation.

For expression-stratified interaction analysis, bins were assigned to expression quintiles based on gene-level transcript abundance. Mean interaction coefficients (from the Poisson regression model) were computed for all pairwise quintile combinations, separately for local (<92 kb) and non-local (≥92 kb) contacts. Enrichment was calculated by dividing each quintile-pair mean by the global mean coefficient across all pairs. Differences between expression quintile groups were assessed using two-sided Mann–Whitney U tests.

